# 3D genome organization during TGFB-induced transcription requires nuclear microRNA and G-quadruplexes

**DOI:** 10.1101/2023.12.22.573061

**Authors:** Julio Cordero, Guruprasadh Swaminathan, Diana G Rogel-Ayala, Karla Rubio, Adel Elsherbiny, Stefan Günther, Thomas Braun, Gergana Dobreva, Guillermo Barreto

## Abstract

Studying the dynamics of three-dimensional (3D) chromatin structure is essential to understand biological processes in the cell nucleus. Recent publications based on integrative analysis of multi-omics studies have provided comprehensive and multilevel insights into 3D genome organization emphasizing its role during transcriptional regulation. While enhancers are regulatory elements that play a central role in the spatiotemporal control of gene expression, chromatin looping has been broadly accepted as a means for enhancer-promoter interactions allowing them to stablish cell-type-specific gene expression signatures. On the other hand, G-quadruplexes (G4s) are non-canonical DNA secondary structures that are both, enriched at promoters and related to increased gene expression. However, the role of G4s in promoter-distal regulatory elements, such as super-enhancers (SE), as well as in 3D genome organization and chromatin looping mediating long-range enhancer-promoter interactions has remained elusive. Here we show that mature microRNA 9 (*miR-9*) is enriched at promoters and SE of genes that are inducible by tissue growth factor beta 1 (TGFB1) signaling. Further, we found that nuclear *miR-9* is required for chromatin features related to increased transcriptional activity, such as broad domains of the euchromatin histone mark H3K4me3 (histone 3 tri-methylated lysine 4) and G4s. Moreover, we show that nuclear *miR-9* is required for promoter-super-enhancer looping. Our study places a nuclear microRNA in the same structural and functional context with G4s and promoter-enhancer interactions during 3D genome organization and transcriptional activation induced by TGFB1 signaling, a critical regulator of proliferation programs in cancer and fibrosis.

## INTRODUCTION

The nuclear genome in eukaryotic cells consists of DNA molecules packaged into thread-like structures known as chromosomes, which are built of chromatin. Thus, chromatin is the physiological template for biological processes in the nucleus of eukaryotic cells. Studying how chromatin is folded inside the cell nucleus and its dynamic three-dimensional (3D) structure is essential to understand these biological processes that comprise transcription, RNA-splicing, -processing, -editing, DNA-replication, -recombination, and -repair. The chromatin is hierarchically organized at different levels including chromosomal territories, compartments, and self-interacting topologically associating domains (TADs) altogether giving rise to a highly dynamic 3D genome organization ^1, 2^. Remarkably, the structure of the genome is intrinsically associated with its function as shown by extensive correlations between chromatin condensation and related gene transcription. For example, chromatin shows condensed regions, referred to as heterochromatin (by convention, transcriptionally “inactive”), and less condensed regions, referred to as euchromatin (transcriptionally “active”). Transcriptional regulation directly corresponds to the mechanisms of how chromatin can be structurally arranged making it accessible for the transcription machinery ^3^. These mechanisms regulating chromatin structure and transcription comprise histone modifications, histone deposition, nucleosome remodeling, DNA methylation, non-coding RNAs (ncRNA), and secondary structures of nucleic acids, among others ^4–9^. In addition, an increasing number of recent publications based on integrative analysis of multi-omics studies implementing next-generation sequencing (NGS) technologies, chromosome conformation capture-based methods, and super-resolution microscopy have provided comprehensive and multilevel insights into 3D genome organization emphasizing its role during transcriptional regulation ^10^.

Chromatin structure alone does not determine the functional status of a gene, but it effectively enables RNA polymerase II (Poll II) recruitment to the promoters, as well as binding of transcription factors, co-activators, co-repressors to DNA sequences that function as regulatory elements controlling gene expression ^11^. A promoter is a sequence of DNA to which proteins bind to initiate transcription of RNA molecules that are usually complementary to the DNA sequence that is located 3’ of the promoter. On the other hand, enhancers are relatively short (∼100–1000 bp) DNA sequences that are bound by transcription factors and regulate gene transcription independent of their distance, location, or orientation relative to their cognate promoter ^12–14^. Super-enhancers (SE) have been proposed to be long genomic domains consisting of clusters of transcriptional enhancers enriched with histone modification markers (such as histone 3 mono-methylated at lysine 4 or acetylated at lysine 27, H3K4me1 and H3K27ac respectively), cofactors (such as mediator of RNA polymerase II transcription subunit 1, MED1, and components of the multimeric protein complex Cohesin), chromatin modifying proteins (such as E1A Binding Protein P300, EP300) and cell-type-specific transcription factors ^15–17^. One versatile feature of chromatin is its ability to form a loop, mediating long-range interactions in which two distant sequences of DNA come to close physical proximity. Chromatin looping has been broadly accepted as a means for enhancer-promoter interactions ^18, 19^.

In addition to the predominant DNA double-helix structure, there are different non-canonical DNA secondary structures, including G-quadruplex (G4), R-loop, H-DNA, Z-DNA, etc. ^6^. A G4 is a stable nucleic acid secondary structure formed by square planes, in which four guanines located in the same plane stabilized by a monovalent cation ^20, 21^. While the early work about G4s mainly focused on their roles in telomeres ^22^, recent studies demonstrated that G4s are both, enriched at promoters ^23, 24^ and related to increased gene expression ^25–29^. On the other hand, G4s were also located in gene bodies and related to reduced gene expression by inhibiting elongation of RNA polymerase ^30, 31^. All these previous studies characterized the biological function of G4s in promoter or promoter-proximal regions enhancing or reducing gene expression depending on the relative position of G4s. Nevertheless, the role of G4s in promoter-distal regulatory elements, such as SE, as well as in chromatin looping mediating long-range enhancer-promoter interactions remains unclear.

The majority of the eukaryotic genome is transcribed into ncRNAs including microRNAs (miRNAs, 21-25 nucleotides long) and long non-coding RNAs (lncRNAs, >200 nucleotides long) ^32^. LncRNAs are important regulators of different biological processes in the nucleus ^33^. Together with other factors, lncRNAs provide a framework for the assembly of defined chromatin structures at specific loci, thereby modulating gene expression, centromere function and silencing of repetitive DNA elements ^33–36^. Although miRNAs are assumed to act primarily in the cytosol by inhibiting translation ^37^, mature miRNAs have been also reported in the nuclei of different cells ^8, 38–41^. Interestingly, a hexanucleotide element has been reported to direct miRNA nuclear import ^42^, nevertheless, the function of miRNAs in the cell nucleus has been sparsely studied. Here we report on microRNA-9 (*miR-9*), which even though its nucleotide sequence is highly conserved across species, also shows high diversity in expression patterns and biological functions depending on the cellular context ^43, 44^. For example, *miR-9* has been reported to target the lncRNA *MALAT1* for degradation in the cell nucleus ^41^. However, it has not been linked to transcription regulation, chromatin structure nor 3D genome organization. Here we propose a mechanism of transcriptional regulation of transforming growth factor beta 1 (TGFB1) responsive genes that requires nuclear *miR-9* and involves G4s and promoter-SE looping.

## RESULTS

### Mature *miR-9* is detected in the cell nucleus enriched at promoters and introns

A phylogenetic tree that was generated using the sequences of mature mouse miRNAs and a heat map comparing their sequence similarity showed that *miR-9* clustered together with miRNAs that have been functionally characterized in the cell nucleus, such as *miR-29b-3p* ^42^, *miR-126-5p* ^45^ and *let-7d-5p* ^8^ (Figure 1a, top). Sequence alignment between mouse *miR-126-5p*, *miR-9* and *miR-29b-3p* showed that various nucleotides are conserved in a sequence stretch that was reported as nuclear shuttling motif from *miR-126-5p* ^45^ (Figure 1a, bottom). We called the partially conserved sequence 5‘-AKYACCWUUUGRUWA-3’ as expanded miRNA nuclear shuttling motif. In addition, we found that the human orthologs of these mature miRNAs, *hsa-miR-126-5P*, *hsa-miR-29B-3P* and *hsa-miR-9-5P*, also contain this expanded miRNA nuclear shuttling motif (Supplementary Figure 1a), showing that it is conserved across species. To confirm the nuclear localization of mature *miR-9*, we performed expression analysis after cell fraction using TaqMan assays specific for mature *miR-9* and total RNA isolated from the cytosolic and the nuclear fractions of different cells (Figure 1b and Supplementary Figure 1b), including mouse lung fibroblasts (MLg and MFML4), mouse lung epithelial cells (MLE-12), mouse mammary gland epithelial cells (NMuMG), and primary human lung fibroblasts (hLF). We detected mature *miR-9* not only in the cytosolic fraction but also in the nuclear fraction of all cells analyzed. Interestingly, the relative levels of nuclear *miR-9* were higher in mouse fibroblasts when compared to epithelial cells. Further, the nuclear localization of *miR-9* was confirmed by RNA fluorescence in situ hybridization (FISH) in MLg cells (Figure 1c, top; Supplementary Figure 1c, left), which also showed that specific regions of the nucleus were *miR-9* depleted. These results were further validated by loss-of-function (LOF) experiments using unlabeled *miR-9*-specific antagomiR probes (Figure 1c, bottom; Supplementary Figure 1c, right). All these results suggest that *miR-9* has a function in the cell nucleus. To investigate the role of *miR-9* in the cell nucleus we performed a sequencing experiment after chromatin isolation by miRNA purification (ChIRP-seq, Figure 1d and Supplementary Figure 1d) using chromatin from MLg cells and control (Ctrl) or *miR-9*-specific biotinylated antisense oligonucleotides for the precipitation of endogenous mature *miR-9* along with the chromatin bound to it. The genome-wide binding profile analysis of *miR-9* (Figure 1d) revealed over twofold increase in the number of *miR-9* peaks at promoters and intronic regions compared to the annotated promoters and intronic regions in the mouse genome. The loci of these putative *miR-9* targets were distributed on all chromosomes (*n* =*2,637*; Supplementary Figure 1e-f). From the *miR-9* ChIRP-seq results we selected putative *miR-9* target genes (*Zdhhc5*, *Ncl*, *Lzts2* and *Hdac7*) for further analysis. Visualization of the loci of these putative *miR-9* target genes using the integrative genomic viewer (IGV) (Figure 1e) showed *miR-9* enrichment at the promoters, whereas no *miR-9* enrichment was detected at the promoter of the negative control *Prap1*. These results were confirmed by quantitative PCR after ChIRP using promoter-specific primers, chromatin from MLg cells and Ctrl or *miR-9*-specific biotinylated antisense oligonucleotides for the precipitation of endogenous *miR-9*-chromatin complexes (Figure 1f). Our results together demonstrate that mature *miR-9* is in the cell nucleus and directly binds to promoters of putative *miR-9*-target genes suggesting a potential role in transcription regulation.

**Figure 1:**
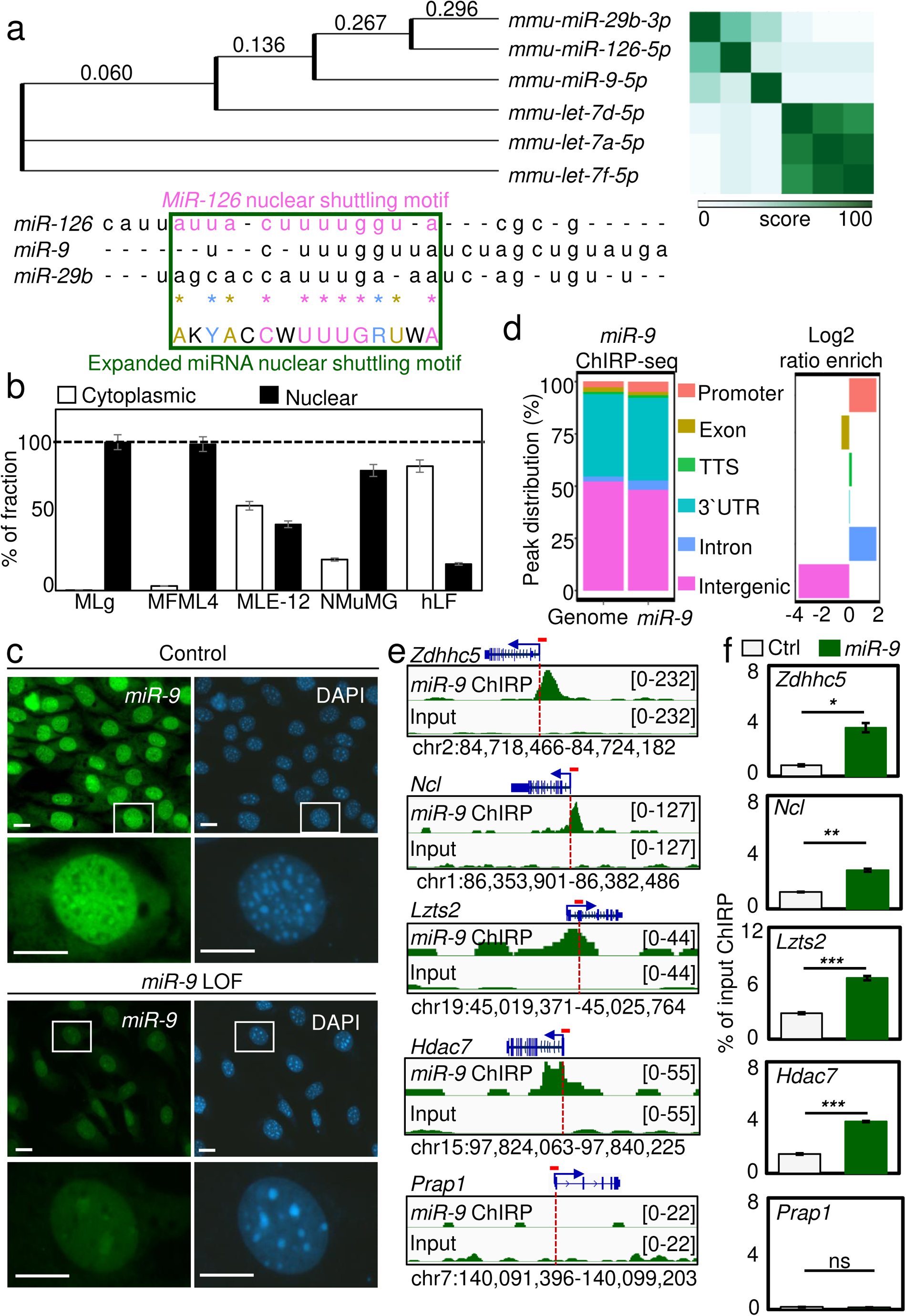
Mature *miR-9* is detected in the cell nucleus enriched at promoters and introns. (**a**) Phylogenetic tree (left) and a heat map (right) that were generated using the sequences of the indicated mature mouse miRNAs. Numbers indicate distance score. Bottom, sequence alignment of the indicated mature mouse miRNAs highlighting the published *miR-126* nuclear shuttling motif (magenta) and the expanded miRNA nuclear shuttling motif (green square) using the IUPAC nucleotide code. Pink letters are conserved among all sequences. Golden letters are conserved in at least 2 sequences. Blue letters indicate conserved type of base (either purine or pyrimidine). (**b**) Mature *miR-9*-specific TaqMan assay after cellular fractionation (Supplementary Figure 1b) of indicated cell lines. Data are shown as means ± s.e.m. (*n* = 3 independent experiments). (**c**) Fluorescence microscopy of MLg cells after RNA FISH confirmed nuclear localization of endogenous *miR-9*. Cells were transiently transfected with control (top) or *miR-9*-specific antagomiR probes (bottom) to induce a *miR-9* loss-of-function (LOF). Representative images from three independent experiments. Squares are shown at higher magnification. DAPI, nucleus. Scale bars, 10 µm. (**d**) Genome-wide distribution of *miR-9* peaks relative to gene structures by ChIRP-seq in MLg cells. Square on right shows *miR-9* enrichment in different gene structure as Log2 ratios. TTS, transcription termination sites; 3’UTR, 3’untraslated regions. (**e**) Visualization of selected *miR-9* target genes using IGV genome browser showing *miR-9* enrichment in MLg cells. ChIRP-seq reads were normalized using reads per kilobase per million (RPKM) measure and are represented as log2 enrichment over their corresponding inputs. Images show the indicated gene loci with their genomic coordinates. Arrows, direction of the genes; blue boxes, exons; red line, regions selected for single gene analysis in f. (**f**) Analysis of selected *miR-9* target genes by ChIRP using chromatin from MLg cells and control (Ctrl) or *miR-9*-specific biotinylated antisense oligonucleotides for the precipitation of endogenous mature *miR-9* along with the chromatin bound to it. Bar plots presenting data as means; error bars, s.e.m (*n* = 3 biologically independent experiments); asterisks, *P*-values after two-tailed t-test, \*\*\**P* ≤ 0.001; \*\**P* ≤ 0.01; \**P* ≤ 0.05. See also Supplementary Figure 1. Source data are provided as a Source Data file 01.

### *MiR-9* is required for H3K4me3 broad domains, high basal transcriptional activity, and G-quadruplex formation at promoters

To further investigate a potential role of nuclear *miR-9* in transcription regulation, we performed a sequencing experiment after chromatin immunoprecipitation assay (ChIP-seq, Figure 2a, Supplementary Figure 2a) using antibodies specific for the euchromatin histone mark, tri-methylated lysine 4 of histone 3 (H3K4me3), and chromatin from MLg cells that were transiently transfected with Ctrl or *miR9*-specific antagomiR to induce a *miR-9*-LOF. Interestingly, *miR-9*-LOF significantly reduced the broad domains of H3K4me3 from 55% in Ctrl transfected cells to 27% after *miR-9*-LOF (*P* < 0.01), whereas medium and narrow H3K4me3 domains increased. The reduction of H3K4me3 broad domains after *miR-9*-LOF was confirmed by confocal microscopy after H3K4me3-specific immunostaining in Ctrl-and *miR-9*-antagomiR transfected MLg cells (Supplementary Figure 2b). Since broad domains of H3K4me3 have been associated with increased transcription elongation ^46^, we analyzed the transcriptome of MLg cells after *miR-9*-LOF by total RNA sequencing (RNA-seq, Figure 2b-c, Supplementary Figure 2c). Remarkably, from the transcripts that were significantly affected after *miR-9*-LOF (*n* = 3,320), only a minority (*n* =881; 26.5%) showed increased expression after *miR-9*-LOF, whereas 73.5% (*n* = 2,439) showed reduced expression with a median of 1.03 log2 RPKM and an interquartile range (IQR) of 1.51 log2 RPKM (*P* = 5.34E-36), when compared to 1.50 log2 RPKM (IQR = 1.81 log2 RPKM) in Ctrl antagomiR transfected cells (Figure 2c, top). The expression of genes without *miR-9*-enrichment (non-targets) as determined by *miR-9* ChIRP-seq (Figure 1d-e and Supplementary Figure 1d) was non-significantly affected after *miR-9*-LOF (Figure 2c, bottom) supporting the specificity of the *miR9*-specific antagomiR. These results indicate that *miR-9* is required for high basal transcriptional activity of its target genes. This interpretation was supported by ChIP-seq in mouse embryonic fibroblasts (MEF) using antibodies specific for total RNA polymerase II (Pol II) and serine 5 phosphorylated Pol II (Pol II S5p, ^9^) showing transcription initiation. Heat maps representing the results of the *miR-9* ChIRP-seq (Figure 2d), Pol II and Pol II S5p ChIP-seq (Figure 2e) revealed that Pol II and Poll II S5p were enriched at the transcription start sites (TSS) of the genes coding for the transcripts that were significantly affected after *miR-9*-LOF (further referred to as *miR-9* target genes). Moreover, genome-wide precision nuclear run-on assay (PRO-seq) ^47^ and global run-on sequencing (GRO-seq) ^48^, both in MEF, showed nascent RNAs at the TSS of the *miR-9* target genes demonstrating their basal transcriptional activity (Figure 2f). Correlating with these results, we also observed at the TSS of the *miR-9* target genes increased chromatin accessibility and increased H3K4me3 levels by assay for transposase-accessible chromatin with sequencing (ATAC-seq) and ChIP-seq, respectively (Figure 2g). To gain further insights into these results, we performed a motif analysis of the *miR-9* target genes and identified significant enrichment of nucleotides motifs with high G content (Figure 2h), as the ones that favor the formation of G4 ^6, 20^. Remarkably, we found similar motifs significantly enriched in loci that form G4 as determined by G4 ChIP-seq. Further, since G4 has been shown to cooperate with transcription factors at gene promoters ^27, 28^, we analyzed publicly available NGS data that were generated implementing different methods for the assessment of G4 formation (Figure 2i). On one hand, we analyzed ChIP-seq data that were generated using an artificial 6.7 kDa G4 probe (G4P) protein, which binds G4s with high affinity and specificity ^24^. On the other hand, we analyzed NGS data that were generated by G4access, which is an antibody-independent method that relies on moderate nuclease digestion of chromatinized DNA ^28^. Remarkably, we found by both approaches enrichment of G4 at the TSS of the *miR-9* target gene (Figure 2i). Our results demonstrate that the TSS of *miR-9* target genes show (1) reduced nucleosome density, (2) increased levels of the euchromatin histone mark H3K4me3, (3) enrichment of G4, *miR-9* and transcription initiating Poll II S5p, and (4) nascent RNAs.

**Figure 2:**
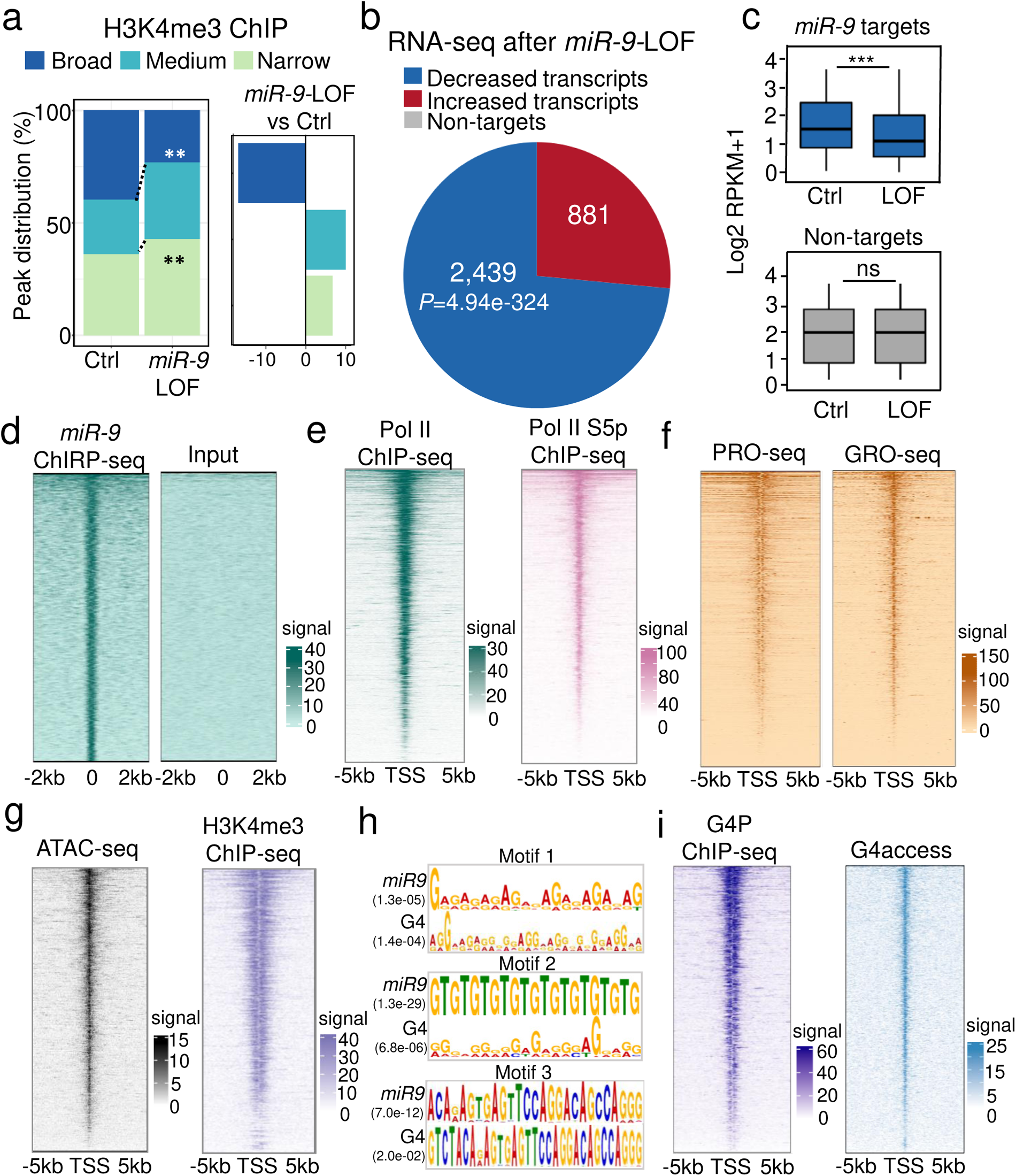
*MiR-9* is required for H3K4me3 broad domains, high basal transcriptional activity, and G-quadruplex formation at promoters. (**a**) Genome-wide distribution of H3K4me3 peaks relative to broad (≥ 2.7 kb), medium (≥ 2 kb and < 2.7 kb) narrow (< 2 kb) H3K4me3 domains in MLg cells that were transiently transfected with control (Ctrl) or *miR-9*-specific antagomiR probes to induce *miR-9* loss-of-function (LOF). Square on right shows H3K4me3 enrichment in different domains as Log2 ratios of MLg cells after *miR-9*-LOF *versus* Ctrl transfected cells. Asterisks, *P*-values after two-tailed t-test, \*\**P* ≤ 0.01. See also Supplementary Figure 2. (**b**) RNA-seq using total RNA from MLg cells that were transfected as in a. Pie chart shows the distribution of transcripts with significantly altered expression levels (*n* = 3,320) between both conditions tested in decreased transcripts (*n* = 2,439) and increased transcripts (*n* = 881) after *miR-9*-LOF. *P*-values after two-tailed t-test. (**c**) Box plots of RNA-seq-based expression analysis of transcripts with significantly decreased levels after *miR-9*-LOF (*miR-9* targets; *n* = 2,439) and with non-significantly (ns) changed levels (non-targets; *n* = 881) after *miR-9*-LOF. Values were normalized using reads per kilobase per million (RPKM) and represented as log2 RPKM. Box plots indicate median (middle line), 25th, 75th percentile (box) and 5th and 95th percentile (whiskers); *P*-values after two-tailed t-test. Source data are provided as Source Data file 01. (**d**) Heat map for *miR-9* enrichment at the TSS ± 2 kb of *miR-9* target genes as determined by RNA-seq in b and c. (**e-g**) Heat maps for enrichment of total Pol II and Pol II S5p (e), nascent RNA by precision nuclear run-on assay (PRO-seq) and global run-on sequencing (GRO-seq) (f), chromatin accessibility by ATAC-seq and H3K4me3 by ChIP-seq (g), at the TSS ± 5 kb of *miR-9* target genes as determined by RNA-seq in b and c. (**h**) Motif analysis of the *miR-9* target genes showed significant enrichment of nucleotide motifs that are similar to the motifs found in loci that form G4 as determined by G4P ChIP-seq. (**i**) Heat maps for G4 enrichment at the TSS ± 5 kb of the *miR-9* target genes by G4P ChIP-seq (left) or G4access (right). See also Supplementary Figure 2. Source data are provided as a Source Data file 01.

Loci visualization of selected *miR-9* target genes (*Zdhhc5*, *Ncl*, *Lzts2*, *Hdac7*, *Gata6* and *Ep300*) using the IGV genome browser (Figure 3a top and Supplementary Figure 3) showed enrichment of the euchromatin histone mark H3K4me3 at the promoters, which was reduced after *miR-9*-LOF. We also observed at the same loci nascent RNA, supporting basal transcriptional activity, and G4 enrichment. Interestingly, a zoom in to the loci revealed G-rich sequences that favor the formation of G4 ^6, 20^ (Figure 3a, bottom, and Supplementary Figure 3). These results were confirmed by promoter analysis of *Zdhhc5*, *Ncl* and *Lzts2* by ChIP using H3K4me3- or G4-specific antibodies and chromatin from MLg cells that were transiently transfected with Ctrl or *miR9*-specific antagomiR probes (Figure 3b). We detected H3K4me3 and G4 enrichment at the promoters of all analyzed *miR-9* target genes in Ctrl antagomiR-transfected cells, which was significantly reduced after *miR-9*-LOF. Our results demonstrate that the promoters of *miR-9* target genes are enriched with H3K4me3 and G4, correlating with the basal transcriptional activity detected by RNA-seq (Figure 2b-c), in a *miR-9*-dependent manner.

**Figure 3:**
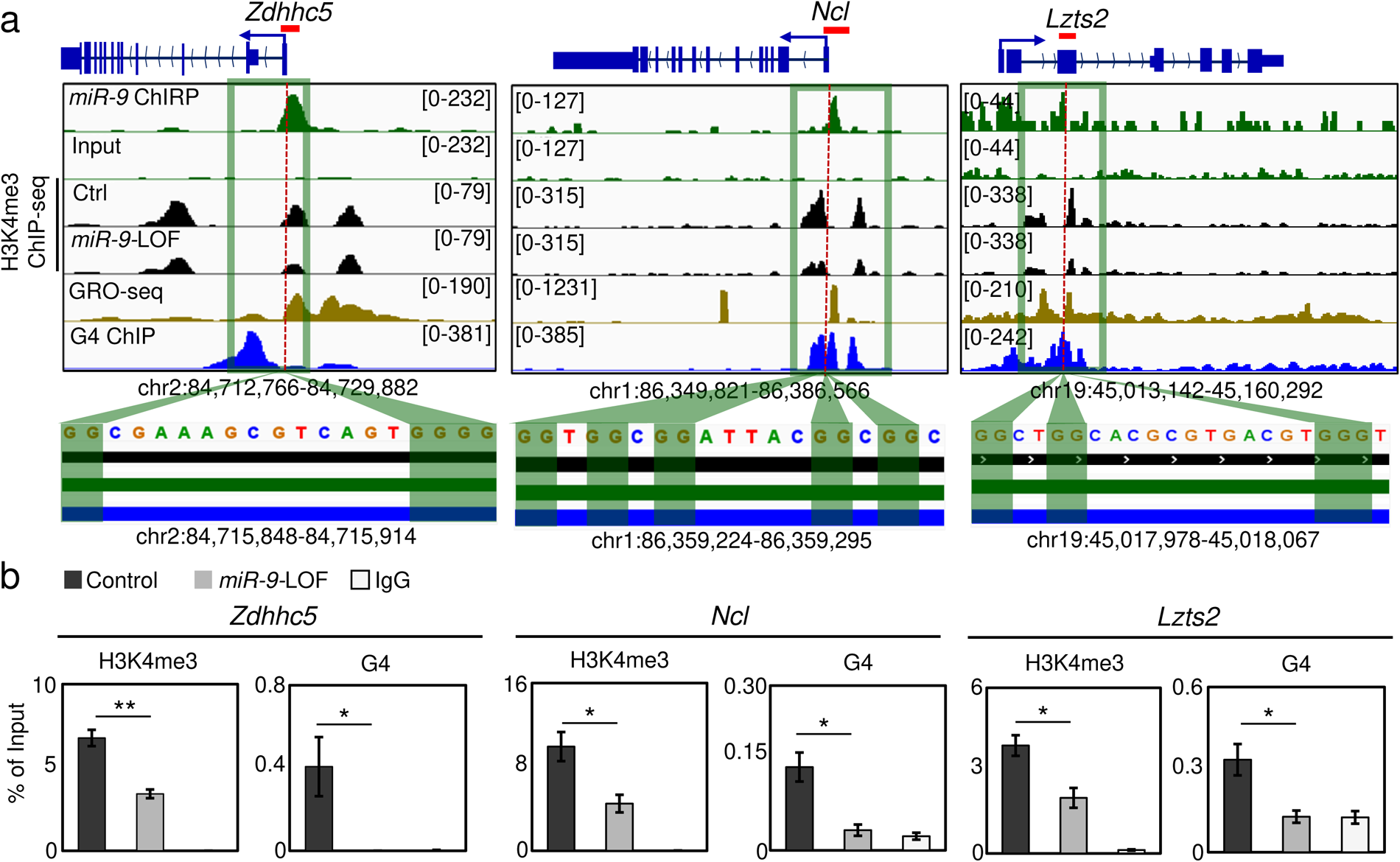
H3K4me3, nascent RNA and G4 are enriched at promoters of selected *miR-9* target genes. (**a**) Visualization of selected *miR-9* target genes using IGV genome browser showing enrichment of *miR-9* by ChIRP-seq (green), H3K4me3 by ChIP-seq in Ctrl and *miR-9*-specifc antagomiR transfected MLg cells (black), nascent RNA by GRO-seq (brown) and G4 by G4P ChIP-seq (blue). Reads were normalized using reads per kilobase per million (RPKM) after bamCoverage. Images show the indicated gene loci with their genomic coordinates. Arrows, direction of the genes; blue boxes, exons; red lines, regions selected for single gene analysis in b; green squares, regions with enrichment of *miR-9*, H3K4me3, nascent RNA and G4; dotted lines, regions shown at the bottom with high G content. (**b**) Analysis of the promoter of selected *miR-9* target genes by ChIP using chromatin from MLg cells transfected with control (Ctrl) or *miR-9*-specific antagomiR to induce *miR-9* loss-of-function (LOF). Bar plots presenting data as means; error bars, s.e.m (*n* = 3 biologically independent experiments); asterisks, *P*-values after two-tailed t-test, \*\**P* ≤ 0.01; \**P* ≤ 0.05. See also Supplementary Figure 3. Source data are provided as a Source Data file 01.

### Nuclear *miR-9* is enriched at super-enhancers and is required for G-quadruplexes

Alluvial plot implementing the data from *miR-9* ChIRP-seq and G4P ChIP-seq ^24^ showed enrichment of *miR-9* and G4 at loci that are also enriched with markers of active SE, such as MED1 and H3K27Ac ^49, 50^ (Figure 4a). Furthermore, a Venn diagram using the same data sets together with the data from a GRO-seq experiment ^48^ showed 3,583 common loci (Figure 4b) suggesting transcriptional activity from these loci. Remarkably, 95.5% (*n* = 3,423) of these transcripts were found in the animal eRNA database ^51^ as enhancer RNAs (eRNA) (Figure 4c). Moreover, mapping the *miR-9* ChIRP-seq data to known loci of typical enhancer (TE) or SE showed that *miR-9* is significantly enriched at SE (Figure 4d). These results were confirmed by further analysis of the *miR-9* ChIRP-seq data together with publicly available ChIP-seq data of proteins that have been related to SE (Figure 4e). Aggregate plots showed enrichment of *miR-9* specifically at SE together with MED1, MYC, H3K27Ac, KLF4, MEIS1, EP300, HDAC1, SMAD3, BRG1, HDAC2, EST1 and RAD21. In addition, we also observed enrichment of CHD4 and SMARCA5 at the same SE as *miR-9*. Supporting these observations, we found that *miR-9* pulled down endogenous CHD4 and SMARCA5 by miRNA pulldown (miR-Pd) followed by Western Blot (Supplementary Figure 4a), whereas both proteins precipitated endogenous *miR-9* by chromatin-RNA immunoprecipitation (Ch-RIP) followed by *miR-9*-specific TaqMan assays (Supplementary Figure 4b). Our results suggest CHD4 and SMARCA5 as components of SE. Remarkably, aggregate plots using G4-specific ChIP-seq data ^24^ showed G4 enrichment at the same SE as *miR-9* in mouse lung fibroblasts (MLg cells) and adenocarcinoma human alveolar basal epithelial cells (A549 cells) (Figure 4f).

**Figure 4:**
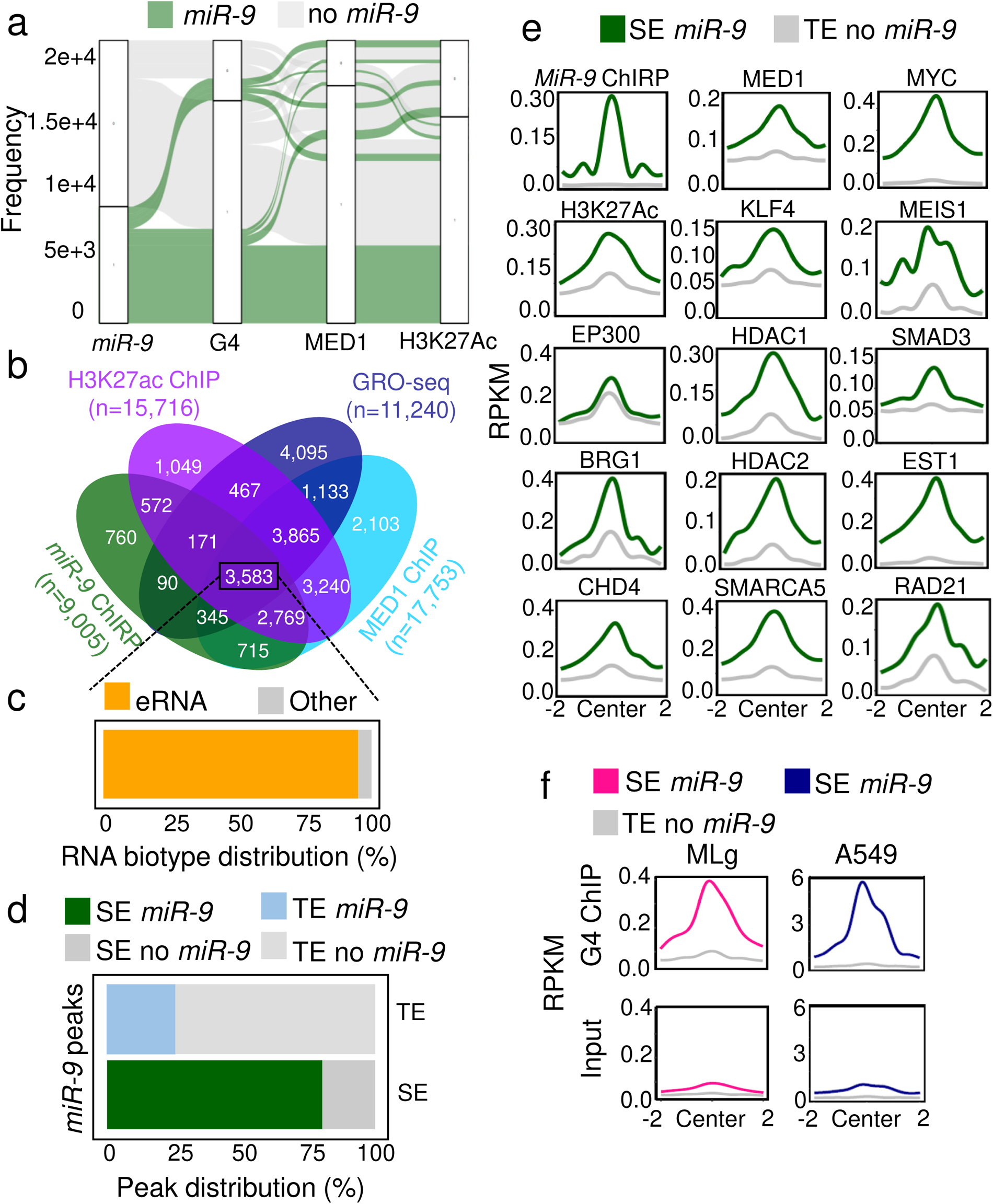
Nuclear *miR-9* is enriched at super-enhancers. (**a**) Alluvial plot showing the proportion of loci with *miR-9* enrichment by ChIRP-seq (green) also with enrichment of G4 by G4P ChIP-seq and the SE markers MED1 and H3K27Ac by ChIP-seq. No-*miR-9*, loci without *miR-9* enrichment. (**b**) Venn diagram showing the number of common loci (*n* = 3,583) with enrichment of *miR-9* by ChIRP-seq (green), H3K27ac (purple) and MED1 (turquoise) by ChIP-seq and nascent RNA (blue) by GRO-seq. (**c**) Venn diagram showing that 3,423 (95.5%) of the transcripts related to the common loci in **b** were found in a database as enhancer RNAs. (**d**) Genome-wide distribution of *miR-9* peaks by ChIRP-seq in MLg cells relative to super-enhancers (SE) and typical enhancers (TE). (**e**) Aggregate plots showing at SE (green lines) and TE (grey lines) ± 2 kb the enrichment of *miR-9* by ChIRP-seq and the indicated proteins by ChIP-seq, which have been previously related to SE in various murine cells. Reads were normalized using reads per kilobase per million (RPKM) measure. (**f**) Aggregate plots showing at SE (magenta and blue lines) and TE (grey lines) ± 2 kb the enrichment of G4 by ChIP-seq in mouse MLg and human A549 cells. Reads were normalized using RPKM. See also Supplementary Figure 4. Source data are provided as a Source Data file 01. Supplementary Table 1 contains all data sets used in this study.

Loci visualization of selected SE with *miR-9* enrichment using the IGV genome browser (Figure 5a top and Supplementary Figure 5) confirmed the enrichment of G4 at the same loci, as well as of MED1, KLF4 and H3K27Ac ^49, 50^, which are markers of active SE. Interestingly, a zoom in to the loci revealed G-rich sequences that favor the formation of G4 ^6, 20^ (Figure 5a, bottom, and Supplementary Figure 5). These results were confirmed by analysis of the loci of these SE by ChIP followed by qPCR using H3K4me3- or G4-specific antibodies and chromatin from MLg cells that were transiently transfected with Ctrl or *miR9*-specific antagomiR probes (Figure 5b). We detected H3K4me3 and G4 enrichment at the loci of all analyzed SE in Ctrl antagomiR-transfected cells, which was significantly reduced after *miR-9*-LOF. Our results demonstrate that SE with *miR-9* enrichment are also enriched with H3K4me3 and G4 in a *miR-9*-dependent manner.

**Figure 5:**
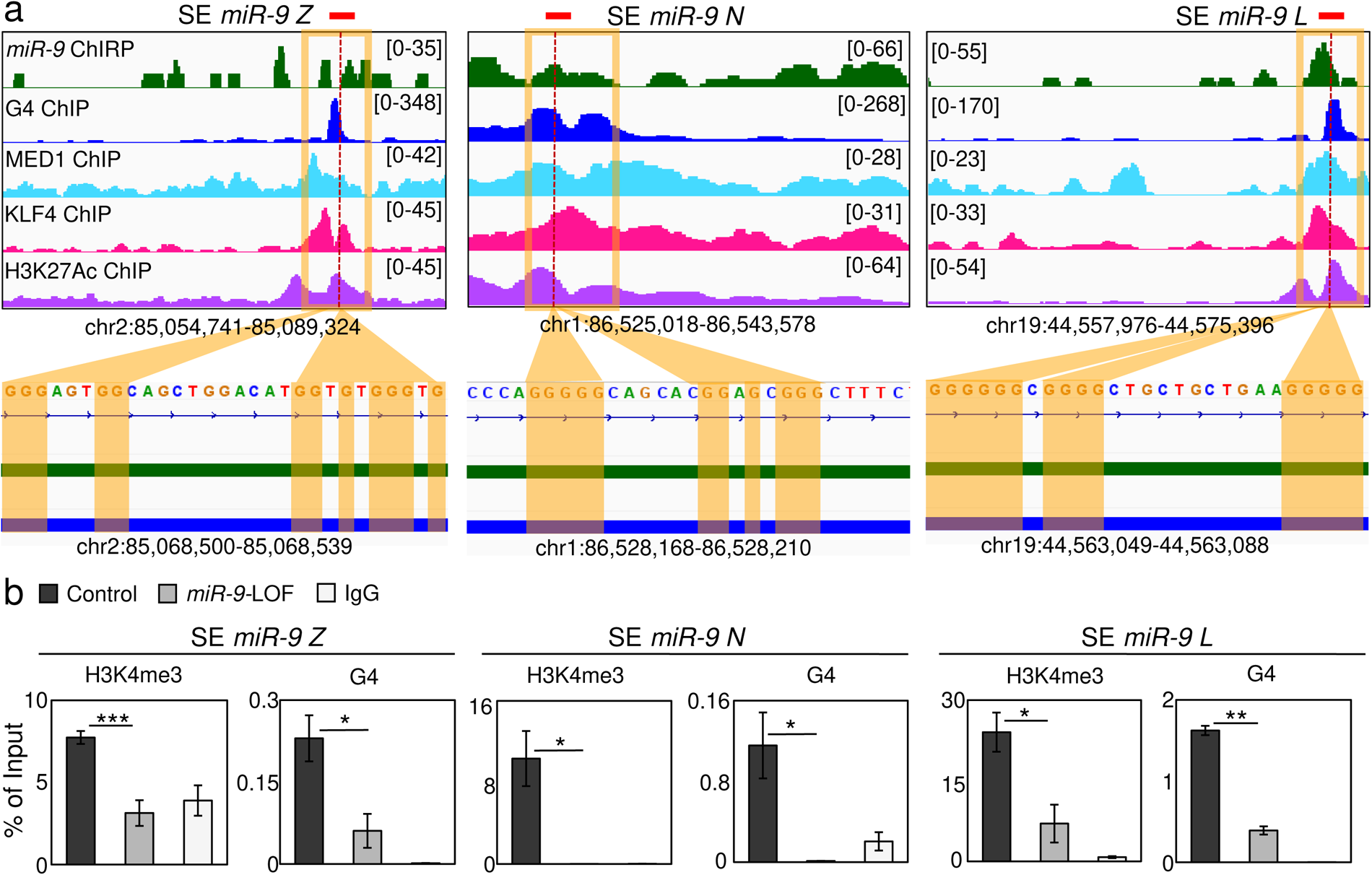
Nuclear *miR-9* is enriched at super-enhancers and is required for G-quadruplexes. (**a**) Visualization of selected SE with *miR-9* enrichment using IGV genome browser showing enrichment *miR-9* by ChIRP-seq (green), G4 by G4P ChIP-seq (blue), MED1 (turquoise), KLF4 (magenta) and H3K27Ac (purple) by ChIP-seq. Reads were normalized using reads per kilobase per million (RPKM). Images show the indicated gene loci with their genomic coordinates. Orange squares, regions with enrichment of *miR-9*, G4 and SE markers; red lines, regions selected for single gene analysis in b; dotted lines, regions shown at the bottom with high G content. (**b**) Analysis of selected SE with *miR-9* enrichment by ChIP using chromatin from MLg cells transfected with control (Ctrl) or *miR-9*-specific antagomiR to induce *miR-9* loss-of-function (LOF). Bar plots presenting data as means; error bars, s.e.m (*n* = 3 biologically independent experiments); asterisks, *P*-values after two-tailed t-test, \*\*\**P* ≤ 0.001; \*\**P* ≤ 0.01; \**P* ≤ 0.05. See also Supplementary Figure 5. Source data are provided as a Source Data file 01.

### Promoter-super-enhancer looping of TGFB1-responsive genes requires *miR-9*

To further elucidate the biological relevance of our findings we performed gene set enrichment analysis (GSEA) ^52^ of the loci with *miR-9* enrichment and nascent RNA as determined by *miR-9* ChIRP-seq and GRO-seq, respectively (Figure 6a-b). We found significant enrichment of genes related to the categories “TGFB cell response” (*P* = 7.55E-09), “TGFB” (*P* = 2.61E-08), “Cell proliferation” (*P* = 4.66E-08) and “Fibroblasts proliferation” (*P* = 1.61E-07), suggesting an involvement of the loci with *miR-9* enrichment and nascent RNA in these biological processes. To link this observation with our results showing reduction of H3K4me3 after *miR-9*-LOF at specific promoters (Figure 3b) and SE (Figure 5b) we did ChIP-seq using H3K4me3-specific antibodies and chromatin from MLg cells that were transiently transfected with Ctrl or *miR9*-specific antagomiR probes, and non-treated or treated with TGFB1 (Figure 6c and Supplementary Figure 6a-b). Loci visualization of the selected *miR-9* target genes (*Zdhhc5*, *Ncl*, *Lzts2*, *Hdac7*, *Gata6* and *Ep300*) using the IGV genome browser showed H3K4me3 enrichment at the promoters in non-treated, and Ctrl antagomiR transfected MLg cells that increased after TGFB1 treatment. However, a combination of TGFB1 treatment and *miR9*-specific antagomiR transfection showed that *miR-9*-LOF counteracted the effect caused by TGFB1 demonstrating the requirement of *miR-9* for the chromatin changes induced by TGFB1 and suggesting its requirement for TGFB1-inducibility of the analyzed genes, as shown below.

**Figure 6:**
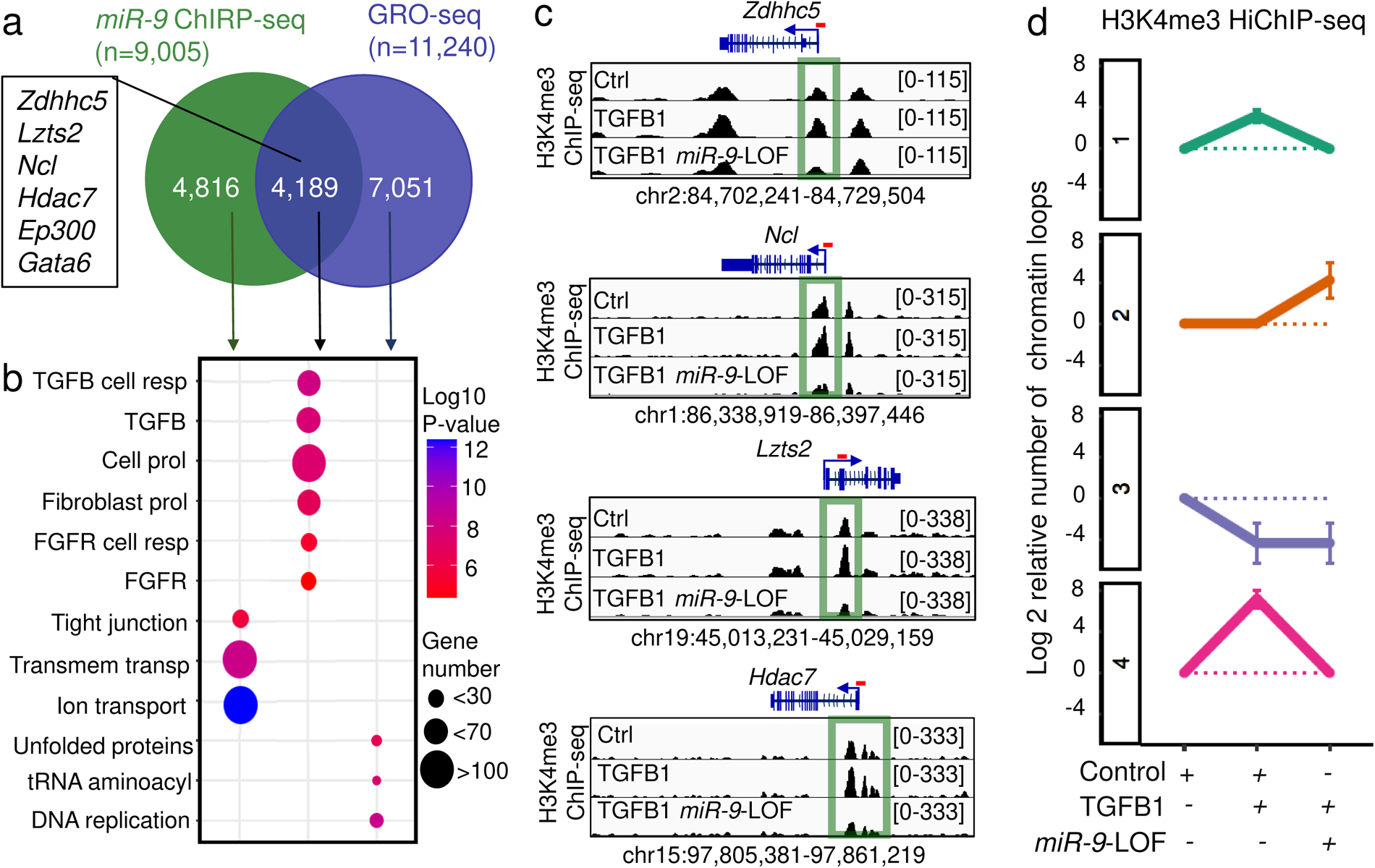
Nuclear *miR-9* is required for H3K4me3 enrichment and chromatin loops at promoters of TGFB1-responsive genes. (**a**) Venn diagram after cross analysis of *miR-9* ChIRP-seq and GRO-seq ^48^ showing loci with *miR-9* enrichment and nascent RNAs (n = 4,189), in which the selected *miR-9* target genes are included (square). (**b**) Gene set enrichment analysis (GSEA) of the three loci groups identified in **a**. Resp, response; prol, proliferation; trans, transport. *P* values after two-tailed Fisheŕs exact test and represented as Log10. (**c**) Visualization of selected *miR-9* target genes using IGV genome browser showing enrichment H3K4me3 by ChIP-seq in MLg cells that were transfected with control (Ctrl) or *miR-9*-specific antagomiR to induce a loss-of-function (LOF), and non-treated or treated with TGFB1, as indicated. Images show the indicated loci with their genomic coordinates. Arrows, transcription direction; green squares, promoter regions; dotted lines, regions selected for single gene analysis in Figure 3b. (**d**) Line charts showing the number of significant chromatin interactions at loci with *miR-9* and H3K4me3 enrichment in MLg cells treated as in **c**. Four clusters were generated using k-means algorithm. Data are represented as Log2 of the ratio relative to Ctrl-transfected, non-treated cells. See also Supplementary Figure 6 and 7. Source data are provided as a Source Data file 01.

Since we observed enrichment of *miR-9*, H3K4me3 and G4 not only at promoters of *miR-9* target genes (Figures 1-3), but also in specific SE (Figures 4-5), we decided to analyze the genome-wide effect of TGFB1 on chromatin conformation by a technique that consists of an *in situ* Hi-C library preparation followed by a chromatin immunoprecipitation (HiChIP). For this HiChIP-seq we used H3K4me3-specific antibodies to precipitate active SE and chromatin from MLg cells that were transiently transfected with Ctrl or *miR9*-specific antagomiR probes, and non-treated or treated with TGFB1 (Figure 6d and Supplementary Figure 7). Analysis of the H3K4me3-specific HiChIP-seq data by k-means clustering revealed four clusters. We focused on clusters 1 and 3 since we observed in these two clusters an increase of chromatin interactions in response to TGFB1 treatment in a *miR-9*-dependent manner, since *miR-9*-LOF reduced the effect caused by TGFB1. Interestingly, in these two clusters we found the loci of the *miR-9* targets (*Zdhhc5*, *Ncl*, *Lzts2* and *Hdac7*) that were selected based on our *miR-9* ChIRP-seq (Figure 1c-e). Further, we generated IGV genome browser snapshots to visualize the enrichment of *miR-9*, G4, MED1, KLF4 and H3K27ac at the loci of promoters of *miR-9* target genes and SE with *miR-9* enrichment (Figure 7a, top). In the same snapshots we presented the results of the H3K4me3-specific HiChIP-seq (Figure 7a, bottom) showing chromatin loops between the promoters of *miR-9* target genes and SE with *miR-9* enrichment in Ctrl antagomiR-transfected cells. Strikingly, these promoter-SE-loops increased in TGFB1 treated cells in a *miR-9*-dependent manner, since *miR-9*-LOF counteracted the effect caused by TGFB1. To correlate the results from H3K4me3-specific HiChIP-seq with changes in chromatin, we analyzed the promoters and SE with *miR-9* enrichment by ChIP qPCR using H3K4me3- and G4-specific antibodies and chromatin from MLg cells that were transfected with Ctrl or *miR-9*-specific antagomiRs, and non-treated or treated with TGFB1 (Figure 7b). TGFB1 treatment significantly increased H3K4me3 and G4 levels at the analyzed promoters and SE. Further, *miR-9*-LOF significantly reduced the effect caused by TGFB1 treatment, whereas *miR-9*-LOF alone significantly reduced H3K4me3 and G4 levels when compared to Ctrl antagomiR transfected cells. To correlate these changes in chromatin structure with gene expression, we analyzed the expression of *miR-9* target genes by qRT-PCR in MLg cells under the same conditions as specified above (Figure 8a). The expression of all analyzed *miR-9* target genes significantly increased after TGFB1 treatment in *miR-9*-dependent manner, correlating with our chromatin structure analysis (Figures 6c and 7b). Further, *miR-9*-LOF alone significantly reduced the basal transcription levels of the analyzed *miR-9* target genes when compared to Ctrl antagomiR transfected cells confirming our RNA-seq results (Figure 2b-c and Supplementary Figure 2c). Summarizing, our results support a model (Figure 8b), in which G4s are formed in a *miR-9*-dependent manner at both, promoters of TGFB1-responsive genes, as well as SE with which these promoters form loops (left). Further, H3K4me3, G4 and promoter-SE looping increased after TGFB1 treatment allowing these two regulatory elements to come to close physical proximity and enhance transcription of the corresponding genes (middle) also in *miR-9*-dependent manner (right).

**Figure 7:**
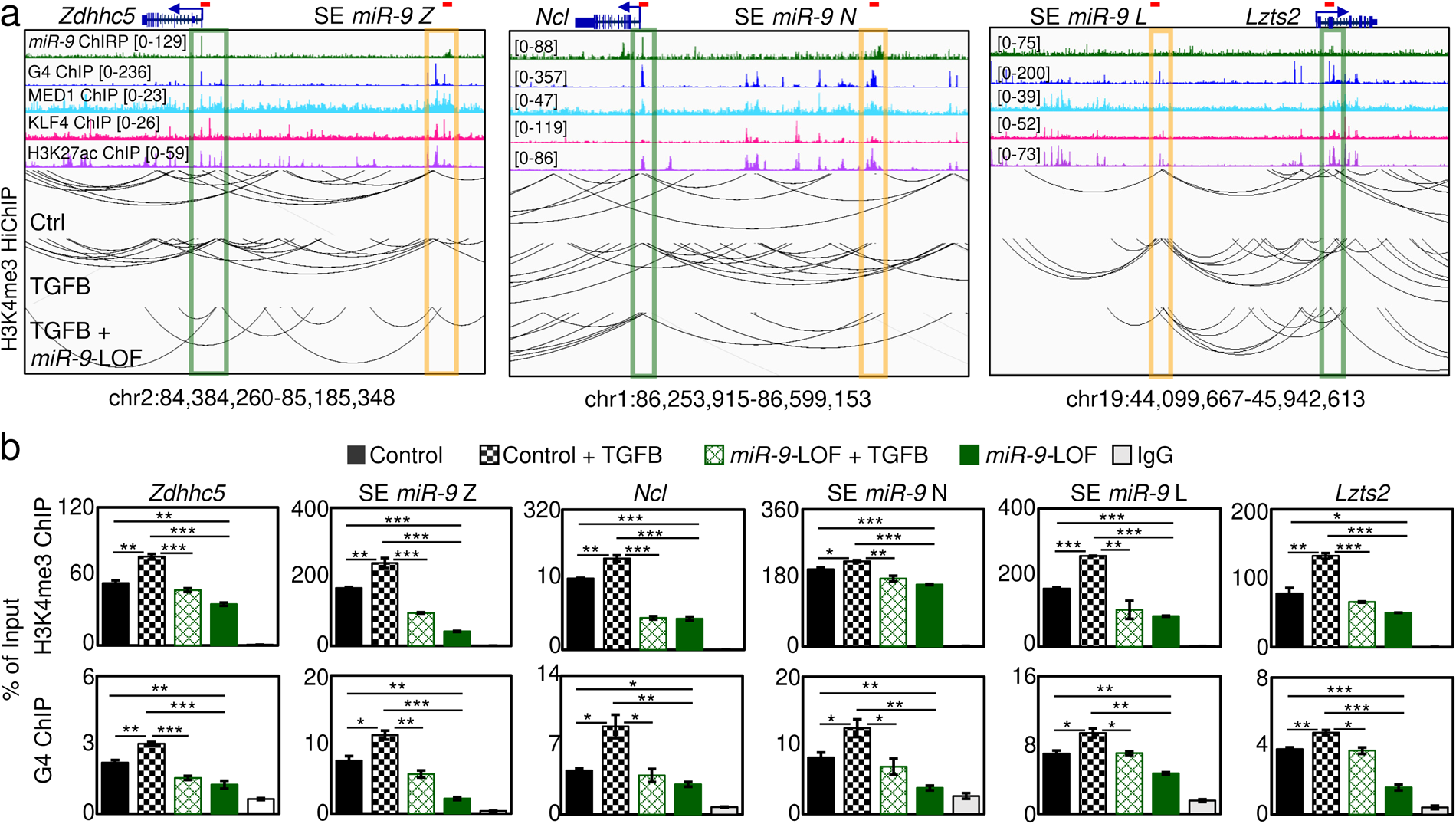
Promoter-super-enhancer looping of TGFB1-responsive genes requires *miR-9*. (**a**) Visualization of promoters of selected *miR-9* target genes (green squares) and SE with *miR-9* enrichment (orange squares) using IGV genome browser showing enrichment *miR-9* by ChIRP-seq (green), G4 by G4P ChIP-seq (blue), MED1 (turquoise), KLF4 (magenta) and H3K27Ac (purple) by ChIP-seq. Reads were normalized using reads per kilobase per million (RPKM) measure and are represented as log2 enrichment over their corresponding inputs. Bottom, chromatin loops by HiChIP-seq in MLg cells that were transfected with control (Ctrl) or *miR-9*-specific antagomir (*miR-9*-LOF, loss-of-function), and non-treated or treated with TGFB1, as indicated. Images show the indicated loci with their genomic coordinates. Arrows, transcription direction; red lines, regions selected for single gene analysis in b. (**b**) Analysis of the promoters and SE highlighted in a by ChIP using the indicated antibodies and chromatin of MLg cells treated as in a. All bar plots present data as means; error bars, s.e.m (*n* = 3 biologically independent experiments); asterisks, *P*-values after two-tailed t-test, \*\*\**P* ≤ 0.001; \*\**P* ≤ 0.01; \**P* ≤ 0.05; ns, non-significant. See also Supplementary Figure 7. Source data are provided as a Source Data file 01.

**Figure 8:**
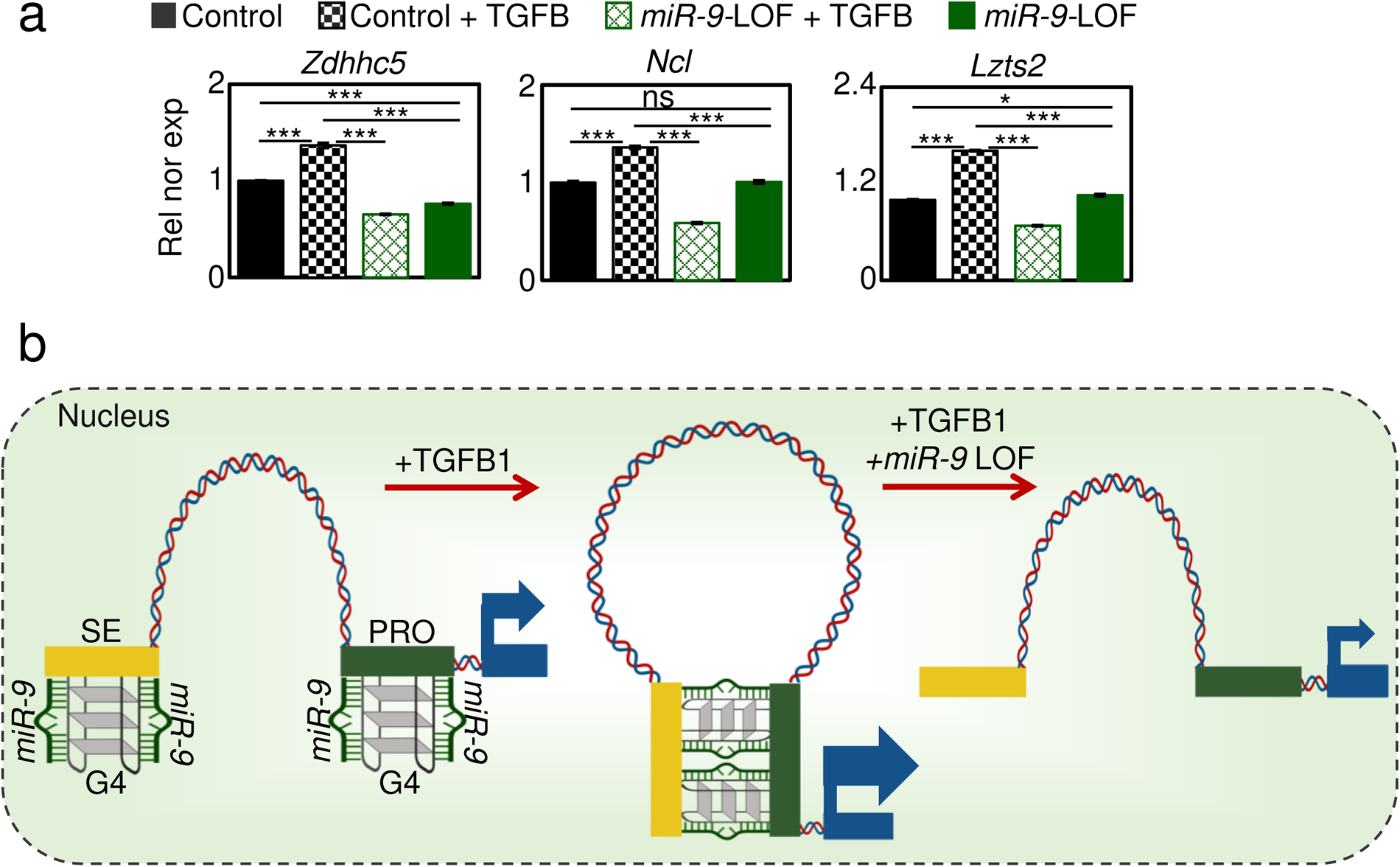
Promoter-super-enhancer looping of TGFB1-responsive genes requires *miR-9*. (**a**) Expression analysis of the selected *miR-9* target genes by qRT-PCR in MLg cells that were transfected with control (Ctrl) or *miR-9*-specific antagomir (*miR-9*-LOF, loss-of-function), and non-treated or treated with TGFB1, as indicated. All bar plots present data as means; error bars, s.e.m (*n* = 3 biologically independent experiments); asterisks, *P*-values after two-tailed t-test, \*\*\**P* ≤ 0.001; \*\**P* ≤ 0.01; \**P* ≤ 0.05; ns, non-significant. See also Supplementary Figure 7. Source data are provided as a Source Data file 01. (**b**) Model summarizing the results presented in the manuscript. Left, G4 are formed in *miR-9* (red lines) -dependent manner at SE (orange box) and promoters (green box) of TGFB1-responsive genes (blue box, coding region). Middle, TGFB1treament increases euchromatin histone mark H3K4me3, G4 and chromatin loops bringing SE and promoter to close physical proximity, thereby enhancing transcription (arrow) of the corresponding gene. Right, *miR-9*-LOF antagonizes the effects induced by TGFB1. Created with BioRender.com

## DISCUSSION

Our study places a nuclear microRNA in the same structural and functional context with non-canonical DNA secondary structures and 3D genome organization during transcription activation. We uncovered a mechanism of transcriptional regulation of TFGB1-responsive genes that requires nuclear *miR-9* and involves G4s and promoter-SE looping. Various aspects of the model proposed here are novel. For example, nuclear *miR-9* had not been related to transcription regulation nor chromatin structure prior to our study, even though *miR-9* participates in a wide spectrum of biological functions including AGO2-dependent degradation of the lncRNA *MALAT1* in the cell nucleus ^41, 43, 44^. Previously, we have shown that other nuclear miRNA, lethal 7d (*Mirlet7d*, also known as *let-7d*), is part of the ncRNA-protein complex MiCEE that mediates epigenetic silencing of bidirectionally transcribed genes and nucleolar organization ^8, 53^. In this context, nuclear *Mirlet7d* binds ncRNAs expressed from these genes, and mediates their degradation by the RNA exosome complex. It will be the scope of future work to determine whether nuclear *miR-9* also binds to ncRNA expressed from the *miR-9* target loci and mediates their degradation by a similar mechanism. Following this line of ideas, it has been reported that G4s are found in genomic regions containing R-loops ^54, 55^, which are three-stranded nucleic acid structures consisting of a DNA-RNA hybrid and the associated non-template single-stranded DNA ^56^. Supporting this line of ideas, we have previously reported that R-loops regulate transcription in response to TGFB1 signaling ^9^. Interestingly, when a G4 is formed opposite to the R-loop on the associated single-stranded DNA, a so called a G-loop structure is generated ^57^. Sato and colleagues recently reported a mechanism involving G-loop structures, in which the transcripts stabilizing the R-loops are relevant for the controlled resolution of the G4s, thereby preventing mutagenic G4s and supporting genomic stability ^57^. We will investigate in a future project whether nuclear *miR-9* is involved in a similar mechanism targeting the ncRNAs in the R-loops and promoting controlled G4 resolution during transcription initiation.

Another interesting aspect of the model proposed here is the participation of G4s in 3D genome organization during transcription activation. DNA G4s are stable four-stranded non-canonical structures that are highly related to promoters and transcription activation ^23–26^. Further, a recent publication based on integrative analysis of multi-omics studies have provided comprehensive mechanistic insights into the function of G4s as promoter elements that reduce nucleosome density, increase the levels of active histone marks (H3K4me3 and H3K27ac), generate nucleosome arrays by positioning nucleosomes at a periodic distance to each other and facilitate pause release of Poll II into effective RNA production resulting in enhanced transcriptional activity ^29^. Even though similar chromatin features were described to be induced by G4s at promoter-distal regulatory elements, such as SE ^58^, no mechanistic insights were provided prior to our study elucidating the role of G4s in chromatin looping mediating long-range enhancer-promoter interactions. We found that G4s are formed in a *miR-9*-dependent manner at both, promoters of TGFB1-responsive genes, as well as SE with which these promoters form loops (Figure 7a-b). Moreover, our results showed promoter-SE looping increase after TGFB1 treatment allowing these two regulatory elements to come to close physical proximity and enhance transcription of the corresponding genes (Figure 8a). One can hypothesize that G4 could stabilize the interaction between these two regulatory elements allowing the increase of transcription. We will investigate this hypothesis in a future project and whether the G4 are formed with DNA strands from the promoter and from the SE.

Our *miR-9*-LOF experiments showed that *miR-9* is required for H3K4me3 broad domains (Figure 2a), basal transcriptional activity (Figure 2b-c) and TGFB1-inducibility (Figure 7c) of *miR-9* target genes. H3K4me3 is a well-characterized euchromatin histone mark related to genes with high transcriptional activity, probably contributing to release of Poll II pausing into elongation ^46, 59^. Furthermore, it has been shown that genes with broad domains of H3K4me3 are transcriptionally more active than genes with narrow domains ^46, 60^, which is consistent with our findings. Interestingly, H3K4me3 broad domains have been linked to genes that are critical to cellular identity and differentiation ^46, 60–62^. Moreover, H3K4me3 broad domains are associated with increased transcription elongation and enhancer activity, which together lead to exceptionally high expression of tumor suppressor genes, including TP53 and PTEN ^46^. On the other hand, TGFB signaling is one of the prominent pathways implicated in hyperproliferative disorders, including cancer ^36, 53, 63–65^. In early stages of cancer, TGFB signaling exhibits tumor suppressive effects by inhibiting cell cycle progression and promoting apoptosis. However, in late stages, TGFB signaling exerts tumor promoting effects, increasing tumor invasiveness, and metastasis. It will be the scope of our future work to determine the clinical relevance of the model of transcription regulation proposed here within the context of hyperproliferative disorders with special attention on carcinogenesis.

## MATERIALS AND METHODS

### Cell culture

Mouse lung fibroblast cells MLg (ATCC CCL-206) and MFML4 ^66^ were cultured in complete DMEM (4.5 g/L glucose, 10% FCS, 1% Penn-strep, 2 mM l-glutamine) at 37°C in 5% CO2. Mouse lung epithelial cells MLE-12 (ATCC CRL-2110) were cultured in complete DMEM/F12 (5% FCS, 1% Penn-strep) at 37^°^C in 5% CO2. Mouse mammary gland epithelial cells NMuMG (ATCC CRL-1636) were cultured in complete DMEM (4.5 g/L glucose, 10mcg/mL insulin (90%), 10% FCS, 1% Penn-strep) at 37 °C in 5% CO_2_. Human primary lung fibroblasts from control donors were cultured in complete MCDB131 medium (8% fetal calf serum (FCS), 1% L-glutamine, penicillin 100 U/ml, streptomycin 0.1 mg/ml, epidermal growth factor 0.5 ng/ml, basic fibroblast growth factor 2 ng/ml, and insulin 5 μg/ml)) at 37°C in 5% CO_2_. During subculturing, cells were 1x PBS washed, trypsinized with 0.25% (w/v) Trypsin and split at the ratio of 1:5 to 1:10. The cell lines used in this paper were mycoplasma free. They were regularly tested for mycoplasma contamination. In addition, they are not listed in the database of commonly misidentified cell lines maintained by ICLAC.

### Cell transfection, treatment and antagomiR-mediated *miR-9* loss-function

Cells were transfected with antagomiR probes (Ambion) using Lipofectamine 2000 (Invitrogen) following the manufacturer’s instructions, and harvested 24 h later for further analysis. Anti-hsa-miR*-Mir9-5p* (Ambion, #17000), and Anti-miR negative control (Ambion, #17010) were transfected at 60 nM final concentrations. Following 20 h after transfection, TGFB1 signaling was induced with 5 ng/ml final concentration of human recombinant TGFB1 (Sigma-Aldrich) for 4 h.

### Bacterial culture and cloning

For cloning experiments, chemically competent *E. coli* TOP10 (ThermoFisher Scientific) were used for plasmid transformation. TOP10 strains were grown in Luria broth (LB) at 37^0^C with shaking at 180rpm for 16h or on LB agar at 37^0^C overnight.

### RNA isolation, reverse transcription, quantitative PCR and TaqMan assay

Expression analysis by qRT-PCR were performed as previously described ^67^. Briefly, total RNA from cell lines was isolated using the RNeasy Mini kit (Qiagen) and quantified using a Nanodrop Spectrophotometer (ThermoFisher Scientific). Synthesis of complementary DNA was performed using 1–2 μg total RNA and the High Capacity cDNA Reverse Transcription kit (Applied Biosystems). Quantitative real-time PCR reactions were performed using SYBR® Green on the Step One plus Real-time PCR system (Applied Biosystems). Housekeeping gene *Gapdh* was used to normalize gene expression. Primer pairs used for gene expression analysis are described in the following table:

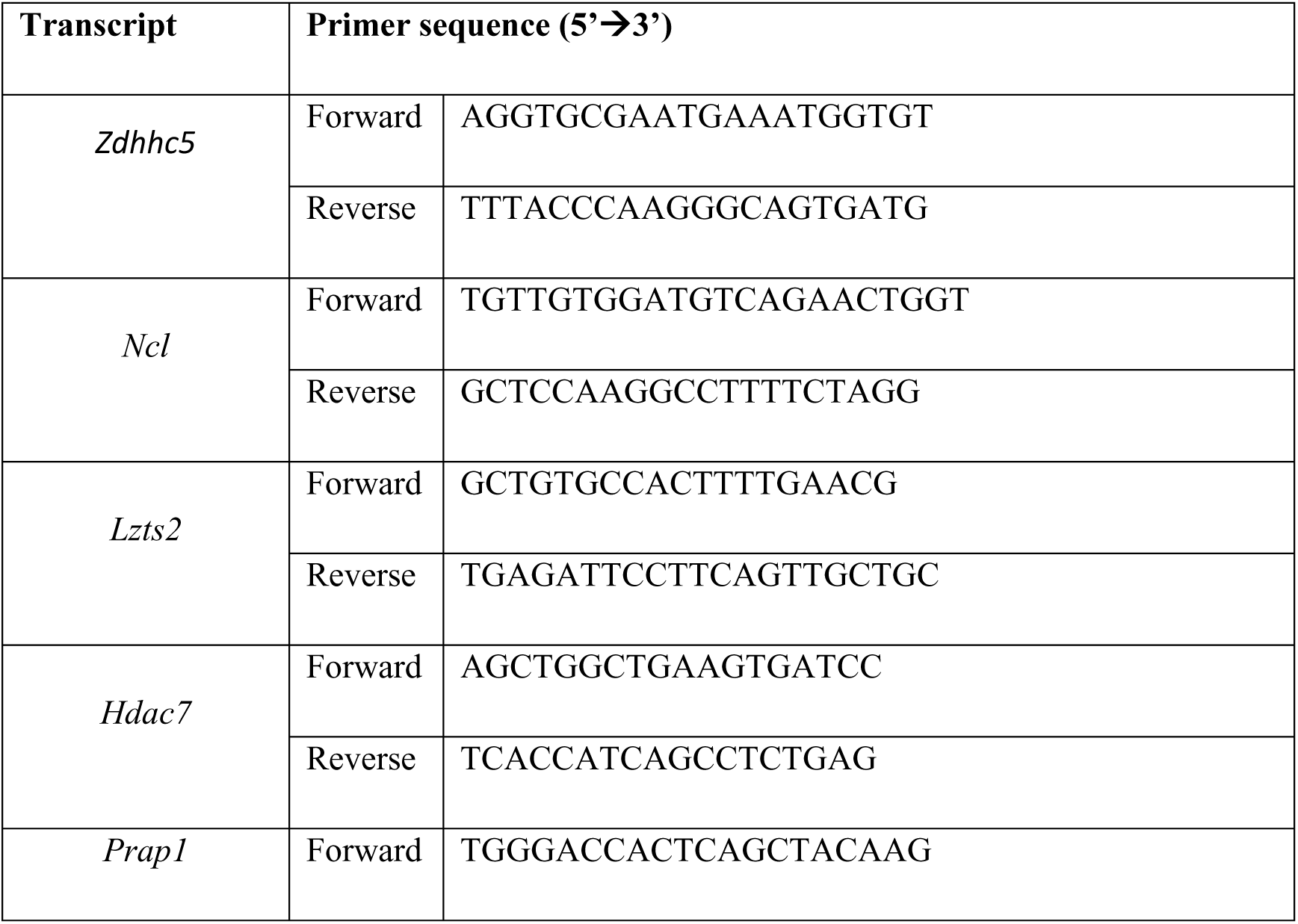
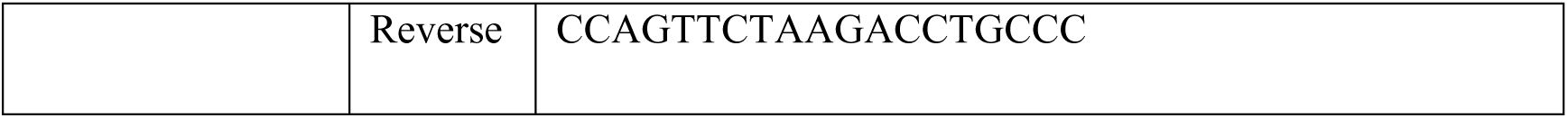

For *miR-9* expression analysis, total RNA was isolated with Trizol (Invitrogen), quantified using a Nanodrop Spectrophotometer (ThermoFisher Scientific), 0.5–2 μg total RNA was used for reverse transcription (High-Capacity cDNA Reverse Transcription Kit, ThermoFisher Scientific) and subsequently *miR-9*-specific TaqMan assay (Applied Biosystems) in the Step One plus Real-time PCR system (Applied Biosystems).

### MiRNA fluorescence in situ hybridization

MiRNA Fluorescence in situ hybridization (MiRNA-FISH) was performed as described earlier ^8^ with minor adaptations. Briefly, cells were fixed with 4% PFA, dehydrated with 70% ethanol and incubated with pre-hybridization buffer (50% formamide, 5X SSC, 5X Denhardt’s solution, 200 μg/ml yeast RNA, 500 μg/ml salmon sperm DNA and 2% Roche blocking reagent in DEPC treated water). Incubation with pre-hybridization buffer was carried out during 4h at room temperature. Pre-hybridization buffer was replaced with denaturizing hybridization buffer (10% CHAPS, 20% Tween, 50% formamide, 5X SSC, 5X Denhardt’s, 200 μg/ml yeast RNA, 500 μg/ml salmon sperm DNA and 2% Roche blocking reagent in DEPC treated water) containing biotin and/or digoxin labeled Locked Nucleic Acid (LNATM) probes (Exiqon) specific to mature *miR-9* and/or miR control respectively to a final concentration of 20 pM and incubated at 55°C overnight. Next day, cells were briefly washed with 5X SSC buffer pre-warmed to 60°C and then incubated with 0.2X SSC at 60°C for 1 h. Later, cells were incubated with B1 solution (0.1 M Tris pH 7.5, 0.15 M NaCl) at room temperature for 10 min. B1 solution was then replaced with blocking solution (10% FCS, 0.1 M Tris pH 7.5, 0.15 M NaCl) and incubated for 1 h at RT. Cells were then incubated with FITC labeled rabbit anti-Biotin (Abcam) antibody overnight at 4°C. DAPI was used as nuclear dye. Cells were examined with a fluorescence microscope (Leica).

### Immunofluorescence and confocal microscopy

Immunostaining was performed as previously described ^53^. Briefly, cells were grown on coverslips, fixed with 4% PFA for 10min at RT and permeabilized with 0.4% Triton-X100/1xPBS for 10min at RT. During immunostaining procedure, all incubations and washes were performed with histobuffer containing 3% bovine serum albumin (BSA) and 0.2% Triton X-100 in 1xPBS, pH7.4. Non-specific binding was blocked by incubating with BSA 5% serum in 1x PBS, pH 7.4. Cells were then incubated with primary antibodies overnight at 4°C. After 3 washes with histobuffer (15 min each), secondary antibody was incubated at room temperature for 1h followed by DAPI nuclear staining (Sigma, Germany). Immunostainings were examined with an immunofluorescence microscope (Leica). Antibodies used were specific anti-H3K4me3 (Abcam, ab8580). Alexa 488 or Alexa 594 tagged secondary antibodies (Invitrogen, Germany, dilution 1:1000) were used. DAPI (Sigma, Germany) was used as nuclear dye.

### Chromatin isolation by miRNA purification and sequencing

Chromatin isolation by miRNA purification (ChIRP) was performed as described ^68^, with slight modifications. Briefly, MLE-12 cells were cross-linked by 1% formaldehyde for 10min, lysed, and sonicated with a Diagenode Bioruptor to disrupt and fragment genomic DNA. After centrifugation, chromatin was incubated with 150 pmol of biotin-labeled anti-sense LNA probes (Exiqon) specific to mature Mirlet7d, or control, at 37 °C for 4h. Streptavidin-magnetic C1 beads (Invitrogen) were blocked with 500ng/μL yeast total RNA and 1mg/mL BSA for 1h at room temperature, and washed three times in nuclear lysis buffer (2mM Tris-HCl, pH 7.0, 250mM NaCl, 2mM EDTA, 2mM EGTA, 1% Triton X-100, 0.2mM DTT, 20mM NaF, 20mM Na3VO4, 40μg/mL phenylmethylsulfonyl fluoride, protease inhibitor, and RNase inhibitor in DEPC-treated water) and resuspended in its original volume. We added 100 µL of washed/blocked C1 beads per 100 pmol of probes, and the whole reaction was mixed for another 30min at 37 °C. Beads:biotinprobes:RNA:chromatin adducts were captured by magnets (Invitrogen) and washed five times with 40×bead volume of wash buffer. DNA was eluted with a cocktail of 100μg/mL RNase A (ThermoFisher Scientific). Chromatin was reverse cross-linked at 65 °C overnight. Later, DNA was purified using the QIAquick PCR Purification Kit (Qiagen) according to manufacturer’s instructions and used for single gene promoter analysis by qPCR and for sequencing. Primer pairs used for qPCR analysis are described in in the sub-section “Chromatin immunoprecipitation” below. ChIRP-seq was performed by single-end sequencing on an Illumina HiSeq2500 machine at the Max Planck-Genome-Centre Cologne. Raw reads were trimmed using Trimmomatic-0.36 with the parameters (ILLUMINACLIP:${ADAPTERS}:2:30:10 LEADING:3 TRAILING:3 SLIDINGWINDOW:4:15 MINLEN:20 CROP:70 HEADCROP:10

(https://doi.org/10.1093/bioinformatics/btu170). Trimmed reads were mapped to mouse genome mm10 using Bowtie2 (default settings) ^69^. Next, PCR duplicates were removed from the BAM files using the MarkDuplicates.jar tool from Picard (version 1.119). MCAS14 ^70^ was used (macs14 -t ChIRP -c Inp -f BAM -p 1e-3 -g 1.87e9 --nomodel --shiftsize 30 -n; and -p 1e-3 -g 1.87e9 --nomodel --shiftsize 100) with the aim to capture genome areas with *miR-9* enrichement. Both peak results were merged with bedtools merge (bedtools merge -d 100 -i cat_PEAKS_s_2.bed) with different annotation with annotatePeaks.pl from homer ^71^. Bam files were converted to bigwig files by the help of bamCoverage from deptools (-bs 20 -- smoothLength 40 -p max --normalizeUsing RPKM -e 150) ^72^. The cis-regulatory element annotation system (CEAS) ^73^ was used to determine the distribution of the peaks from *miR-9* ChIRP-seq in different genomic areas (intron, exon, 5′ UTR, 3′ UTR, promoter, intergenic, and downstream).

### Motif analysis of ChIRP and G4-ChIP-seq data

MEME Suite (Motif-based sequence analysis tools) ^74^ was used for de-novo DNA motif. search analysis. The *miR-9* ChIRP-seq data file containing peaks annotated near the promoter of the genes was used for the motif search. The G4 ChIP-seq ^24^ file containing peaks annotated near the promoter of the genes was used for the motif search was used for the motif search analysis. The settings in the de-novo DNA motifs search were: (1) a normal enrichment mode to search the motif, (2) length of the motifs allow between 10 and 25bp widths, (3) motif site distribution of any time of repetition on the fasta file and (4) three maximum motifs to report.

### Enhancer areas enriched with *miR-9*

To determine the potential super enhancer marked or not marked by *miR-9*, we crossed the *miR-9* ChiRP-seq peaks list with the results from the peak list from H3K27ac peaks from (GSM2417093) determined using the following parameters: findPeaks H3K27ac_maketaglib -i Inp_makeTaglib -region -size 500 -minDist 20000 -fdr 0.05 -F 2 -style super -typical output_enh -superSlope -1000 > OUPUT}_SE.txt.

Those significant SE with more than 100 normalized reads were further crossed with the *miR-9* ChIRP-seq peak by the help of bedtools intersecpt (setings -wa). Those H3K27ac peaks not overlapping the SE list and not overlapping the *miR-9* peaks were considered TE. All ChiP-seq and miR-9 ChIRP-seq were quantified from the center of both Peak list by the help of annotatePeaks from homer with the settings: annotatePeaks.pl list.bed mm10 -size 4000 -norm 1000000 -hist 10 -d maketagLib_Samples > output_quanty.txt. The results of this command was used as input in a custom R-script to produce the aggregate plots (https://github.com/jcorder316/001MIr9_G4_3D_STRUC).

### Chromatin immunoprecipitation (ChIP)

ChIP analysis was performed as described earlier ^9, 75^ with minor adaptations. Briefly, cells were cross-linked with 1% methanol-free formaldehyde (ThermoFisher Scientific) lysed, and sonicated with Diagenode Bioruptor to an average DNA length of 300–600 bp. After centrifugation, the soluble chromatin was immunoprecipitated with 3 µg of antibodies specific for anti-H3K4me3 (Abcam, # ab8580), Anti-DNA G-quadruplex structures Antibody, clone BG4 (Millipore, # MABE917), and anti-IgG (Santa Cruz, #sc-2027). Reverse crosslinked immunoprecipitated chromatin was purified using the QIAquick PCR purification kit (Qiagen) and subjected to ChIP-quantitative PCR. The primer pairs used for gene promoter and super-enhancer regions are described in the following table:

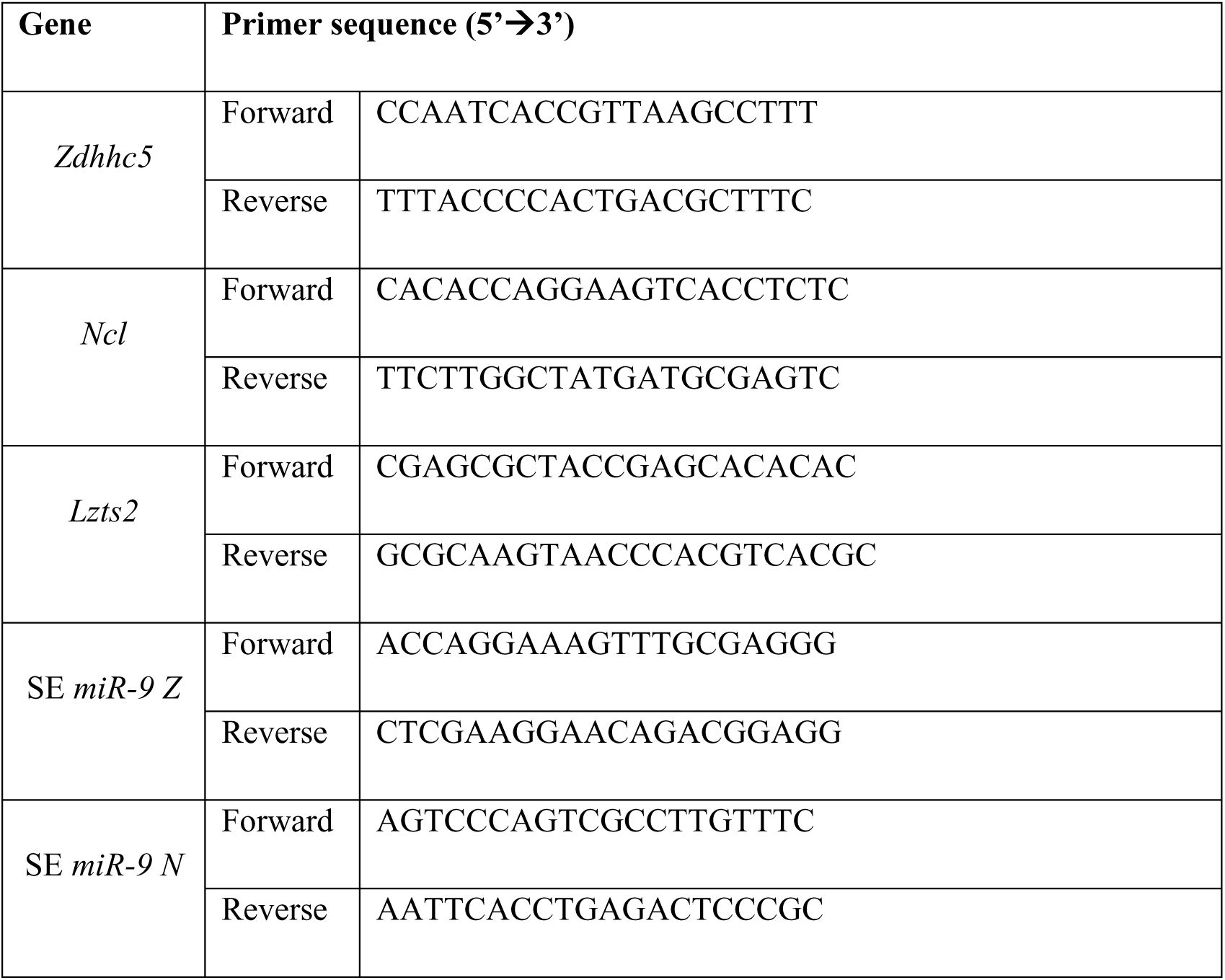
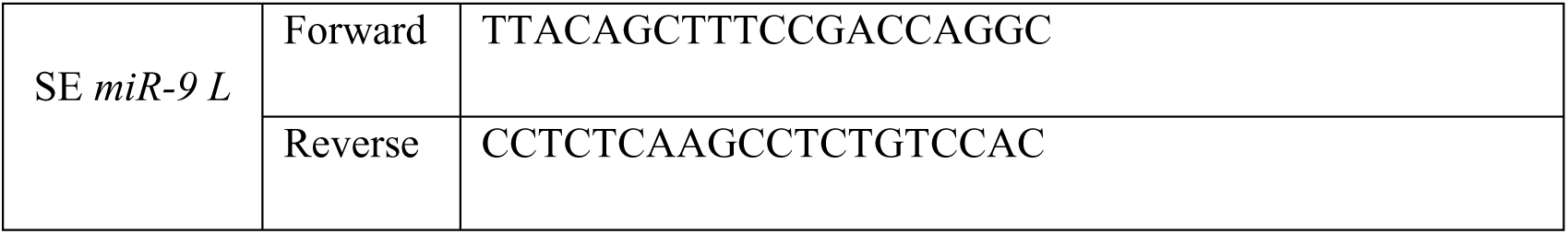

### Chromatin conformation analysis by *in situ* Hi-C library preparation followed by chromatin immunoprecipitation (HiChIP)

HiChIP experiments were performed as previously described ^76^ using antibodies specific for H3K4me3 (Abcam, # ab8580) with the following optimizations: 5-10 million cells were crosslinked with 1% formaldehyde for 10 min at room temperature; prior to restriction digestion, sodium dodecyl sulfate treatment at 62°C for 10 min; restriction digestion with *MboI* (New England Biolabs France, R0147M) for 2 h at 37°C; prior to fill-in reaction, heat inactivation of MboI at 62°C for 10 min followed by 2 washing steps of pelleted nuclei with 1x fill-in reaction buffer; after fill-in reaction, ligation at 4°C for 16 h.

### ChIP sequencing, HiChIP sequencing and data analysis

HiChIP-seq paired-end reads were aligned to the mm10 genome, duplicate reads were removed, reads were assigned to MboI restriction fragments, filtered into valid interactions, and the interaction matrices were generated using the HiC-Pro pipeline default settings ^77^. The contig file of the HiC-Pro was set to allow validPairs at any distance from each other. HiC-Pro valid interaction reads were then used to detect significant interactions using: (1) Maketag libraries were made as followed (makeTagDirectory output_PCA_ucsc smaple_R1_mm10.bwt2merged.bam,sample_R2_mm10.bwt2merged.bam -tbp 1 -genome mm10 -checkGC -restrictionSite GATC. (2) We used (runHiCpca.pl sample_mer25_50 sample_PCA_ucsc -res 25000 -superRes 50000 -genome mm10 -cpu 16) to find the Principal component analysis (PCA) of the data. (3) We used analyzeHiC as followed (analyzeHiC sample_PCA_ucsc -res 1000000 -interactions sample_significantInteractions.txt -nomatrix ).

Only Interaction looping between the H3K4me3 peaks and the TSS (+/-2kb) from *miR-9* candidate genes were considered. The settings were (annotateInteractions.pl Sample_significan mm10 Sample_output_filterk4m9 -filter H3k4me3_HICHIP-seq_peaks -filter2 TSS2kb_fMir9_genes -cpu 16 -washu -pvalue 0.01). (4) The mapped merge bam files ouput from HiC-pro was processed as ChiP-seq to perform Peak calling, using (peak_call -i bam_file -o ouput_peak -r MboI_mm10.txt -f 0.01 -a mm10_chr_size -w 8) from (https://github.com/ChenfuShi/HiChIP_peaks).

To calculate the broadness of H3K4me3 peaks, we first performed summary statistics of the peak size from the merged peak list. If the size of a peak was equal to or higher than the top 75 % of peaks quantile 3 (Q3) was consider wide (≥ 2.7kb). If the peak was between Q3 and Q2 was consider medium size peak (≥ 2 kb and < 2.7 kb). The narrow Hek4me3 peaks were in the bottom 25% peaks, Q1 or less (< 2kb).

### Meta-analysis of ChIP-seq, PRO-seq and GRO-seq

All published data from ChIP-seq, PRO-seq, GRO-seq and ATAC-seq used in this manuscript were listed in the Supplementary Table 1. All these NGS data were downloaded and processed according to the description in their respective publications. Briefly, PRO-seq, GRO-seq and ChIP-seq were processed according to (https://doi.org/10.1038/s41588-018-0139-3). ATAC-seq, G4 ChIP-seq and G4access were processed using Bowtie2 (default settings) and all mapped bam files were converted to bigwig files using the same parameters as for the ChIRP-seq mentioned above.

The enrichment of the factors and nascent RNAs (PRO-seq and GRO-seq) on the TSS of the Down-regularted genes after *miR-9*-LOF were performed by the help of the library from R profileplyr (https://www.bioconductor.org/packages/devel/bioc/vignettes/profileplyr/inst/doc/profileplyr.html) adapting to a custom R-script (https://github.com/jcorder316/001MIr9_G4_3D_STRUC).

### MicroRNA pulldown

For microRNA pulldown (miR-pd), MLE-12 cells were transfected with Biotinylated *mmu-miR-9-5p*-RNA (Exiqon) or Biotinylated *Mirctrl* (Exiqon), at a final concentration of 20 nM. After 48 hours, cells were fixed with 1% glutaraldehyde (Sigma-Aldrich) to preserve RNA-Chromatin interactions. After washing three times with PBS, cell pellets were resuspended in 2 ml hypotonic cell lysis buffer (10 mM Tris-HCl (pH 7.4), 1.5 mM MgCl2, 10 mM KCl, 1 mM DTT, 25 mM NaF, 0.5 mM Na3VO4, 40 μg/ml phenylmethylsulfonyl fluoride, protease inhibitor (Calbiochem) and RNase inhibitor (Promega) in DEPC treated water) on ice for 10 min and then spun down at 700 g for 10 min at 4°C. Nuclear pellets were resuspended in 300 μL of nuclear lysis buffer (50 mM Tris-HCl (pH 7.4), 170 mM NaCl, 20% glycerol, 15 mM EDTA, 0.1% (v/v) Triton X-100, 0.2 mM DTT, 20 mM NaF, 20 mM Na3VO4, 40 μg/ml phenylmethylsulfonyl fluoride, protease inhibitor, RNase inhibitor in DEPC treated water). Nuclear fraction was sonicated (Bioruptor NextGen, Diagenode) at high amplitude for 10 cycles of 30 sec (on/off) pulses. M280 streptavidin magnetic beads (ThermoFisher Scientific) were blocked for 1 h at 4°C in blocking buffer (10 mM Tris-HCl pH 6.5, 1 mM EDTA, and 1 mg/ml BSA) and washed twice with 1 ml washing buffer (10 mM Tris-HCl (pH 7.0), 1 mM EDTA, 0.5 M NaCl, 0.1% (v/v) Triton X-100, RNAase inhibitor and protease inhibitor). Beads were resuspended in 0.5 ml washing buffer. The nuclear extract was then added to the beads and incubated for 1 h at 4°C with slow rotation. The beads were then washed five times with 1 ml washing buffer. RNA bound to the beads (pulled down RNA) and from 10% of the extract (input RNA) was isolated using Trizol reagent LS (Invitrogen) after DNase 1 (NEB) treatment.

For miRNA-protein interaction analysis, nuclear protein extract after DNase1 treatment incubated with 500 pMol Biotinylated *miR-9* or Biotinylated *Mirctrl* for 1 h with shaking at 4°C. M280 streptavidin magnetic beads (ThermoFisher Scientific) were used for pulldown. Protein bound to the beads (miR-Pd), was incubated with 30 μl 2x SDS samples loading buffer, boiled at 95°C for 5min, spun down and loaded on SDS-PAGE for western blot analysis and/or processed for proteomic analysis by mass spectrometry.

### Chromatin RNA immunoprecipitation

Chromatin RNA immunoprecipitation (Ch-RIP) analysis was performed as described ^8^ with minor adaptations. Briefly, cells were cross-linked by 1% formaldehyde for 10 min, lysed, and sonicated with Diagenode Bioruptor to disrupt and fragment genomic DNA. After centrifugation, the soluble chromatin was immunoprecipitated using antibodies. Precipitated chromatin complexes were removed from the beads by incubating with 50 μl of 1% SDS with 0.1 M NaHCO3 for 15 min, vortexing every 5 min and followed by treatment with DNase1.

The collected material was immunoprecipitated using IgG or an HA-specific antibody (Santa Cruz, CatNo. 805). Reverse cross-linked immunoprecipitated materials were used for RNA isolation by Trizol (Invitrogen). Isolated RNA was subjected to cDNA synthesis and further for TaqMan assay specific for mature *miR-9* (Applied Biosystems, Cat. No. 4427975).

### Western blot

Nuclear protein extracts from MLg cells were prepared by cell lysis and nuclei isolation. Briefly, MLg cells were spooled down and washed with PBS. cell pellets were resuspended in 2 ml hypotonic cell lysis buffer (10 mM Tris-HCl (pH 7.4), 1.5 mM MgCl2, 10 mM KCl, 1 mM DTT, 25 mM NaF, 0.5 mM Na3VO4, 40 μg/ml phenylmethylsulfonyl fluoride, protease inhibitor (Calbiochem) and RNase inhibitor (Promega) in DEPC treated water) on ice for 10 min and then spun down at 700 g for 10 min at 4°C. Nuclear pellets were resuspended in 300 μL of nuclear lysis buffer (50 mM Tris-HCl (pH 7.4), 170 mM NaCl, 20% glycerol, 15 mM EDTA, 0.1% (v/v) Triton X-100, 0.2 mM DTT, 20 mM NaF, 20 mM Na3VO4, 40 μg/ml phenylmethylsulfonyl fluoride, protease inhibitor, RNase inhibitor in DEPC treated water). Nuclear fraction was sonicated (Bioruptor NextGen, Diagenode) at high amplitude for 10 cycles of 30 sec (on/off) pulses. Detergent-insoluble material was precipitated by centrifugation at 14,000 rpm for 30 min at 4 °C. The supernatant was transferred to a fresh tube and stored at -20 °C. Protein concentration was estimated using Bradford assay, using serum albumin as standard. 5 μl of serial dilutions of standard protein and samples were mixed with 250 μl of Bradford reagent (500-0205, BIO-RAD Quick Start™).

Western blotting was performed using standard methods and antibodies specific for SMARCA5 (Invitrogen, MA5-35378), LMNB1 (Santa Cruz, sc-374015), CHD4 (Abcam, 70469), RAD21 (Abcam, ab992), GAPDH (Sigma, MFCD01322099). Immunoreactive proteins were visualized with the corresponding HRP-conjugated secondary antibodies (Santa Cruz) using the Super Signal West Femto detection solutions (ThermoFisher Scientific). Signals were detected and analyzed with Luminescent Image Analyzer (Las 4000, Fujifilm).

### RNA sequencing and data analysis

RNA sequencing data for this paper were generated as previously described ^8, 53^. Briefly, total RNA from MLg cells that were transfected with control or *miR-9*-specific antagomiR was isolated using the Trizol method. RNA was treated with DNase (DNase-Free DNase Set, Qiagen) and repurified using the RNeasy micro plus Kit (Qiagen). Total RNA and library integrity were verified on LabChip Gx Touch 24 (Perkin Elmer). Sequencing was performed on the NextSeq500 instrument (Illumina) using v2 chemistry with 1x75bp single end setup. Raw reads were visualized by FastQC to determine the quality of the sequencing. Trimming was performed using trimmomatic with the following parameters: LEADING:3 TRAILING:3 SLIDINGWINDOW:4:15 MINLEN:15 CROP:60 HEADCROP:15. High quality reads were mapped to mm10 uscs genome using with bowtie2 , settings (-D 15 -R 3 -N 0 -L 20 -i S,1,0.56 -k 1). Tag libraries were obtained with MakeTaglibrary from HOMER (default setting). Samples were quantified by using analyzeRepeats.pl with the parameters: mm10 –count exons –strand both –noad. Gene expression was quantified in reads per kilo base million (RPKM) by the help of rpkm.default from EdgeR. Down-regulated genes after *miR-9*-LOF were those genes with a Log2fc (LOF/Ctr)<=-0.58 and Up-regulated those genes with a log2FC >= 0.58.

### Statistical Analysis

Depending on the data, different tests were performed to determine the statistical significance of the results. The values of the statistical tests used in the different experiments can be found in Source Data file 01. Further details of statistical analysis in different experiments are included in the figures and figure legends. Briefly, one set of ChIRP, RNA, ChIP and HiChIP samples were analyzed by deep sequencing. For the rest of the experiments presented here, samples were analyzed at least in triplicates and experiments were performed three times. Statistical analysis was performed using Excel Solver and Prism9. Data in bar plots are represented as mean ± standard error (mean ± s.e.m.). Two-tailed t-tests were used to determine the levels of difference between the groups and *P*-values for significance. *P*-values after two-tailed t-test, \**P* ≤ 0.05; \*\**P* < 0.01, and \*\*\**P* < 0.001.

## Supporting information

Cordero et al 2023 Supplementary Information

## Data availability

The data that support this study are available from the corresponding author upon reasonable request. The source data are provided with this paper as a Source Data file 01. In addition, sequencing data of ChIRP, HiChIP and RNA have been deposited in NCBI’s Gene Expression Omnibus ^78^ and is accessible through SRA Sequence Read Archives NCBI with accession number GSE244952. In addition, we retrieved and used publicly available datasets to aid analysis of our data. Supplementary Table 1 contains all data sets used in this study. The model in Figure 8b was Created with BioRender.com.

## Acknowledgments

We thank Roswitha Bender for technical support; Kerstin Richter and Alessandro Ianni for antibodies; Domenique Dumas for support with confocal microscopy; Amine Armich, Sylvie Fournel-Gigleux, Mohamed Ouzzine, Jean-Baptiste Vincourt, Sandrine Gulberti, Catherine Bui, Lydia Barré and Nick Ramalanjaona for comments.

## Authors Contributions

JC, GS, DGRA, KR, AE, SG and GB designed and performed the experiments; TB and GD were involved in study design; GB and JC designed the study; JC, GB, GS, DGRA, KR and SG analyzed the data; GB, JC, GS and DGRA wrote the manuscript. All authors discussed the results and commented on the manuscript.

