## Supplementary material for "3D genome organization during TGFB-induced transcription requires nuclear microRNA and G-quadruplexes": Cordero et al 2023 Supplementary Information

**This PDF file includes:**

Supplementary Figures 1 to 7

**Other Supplementary Materials for this manuscript include the following:**

Source Data file 01 - This is an Excel file that contains the data for all the plots presented in the article, including the values for statistical significance and the implemented statistical tests.

Supplementary Table 1 – This is an Excel file that contains a list with the accession numbers of all published NGS data sets used in this manuscript.

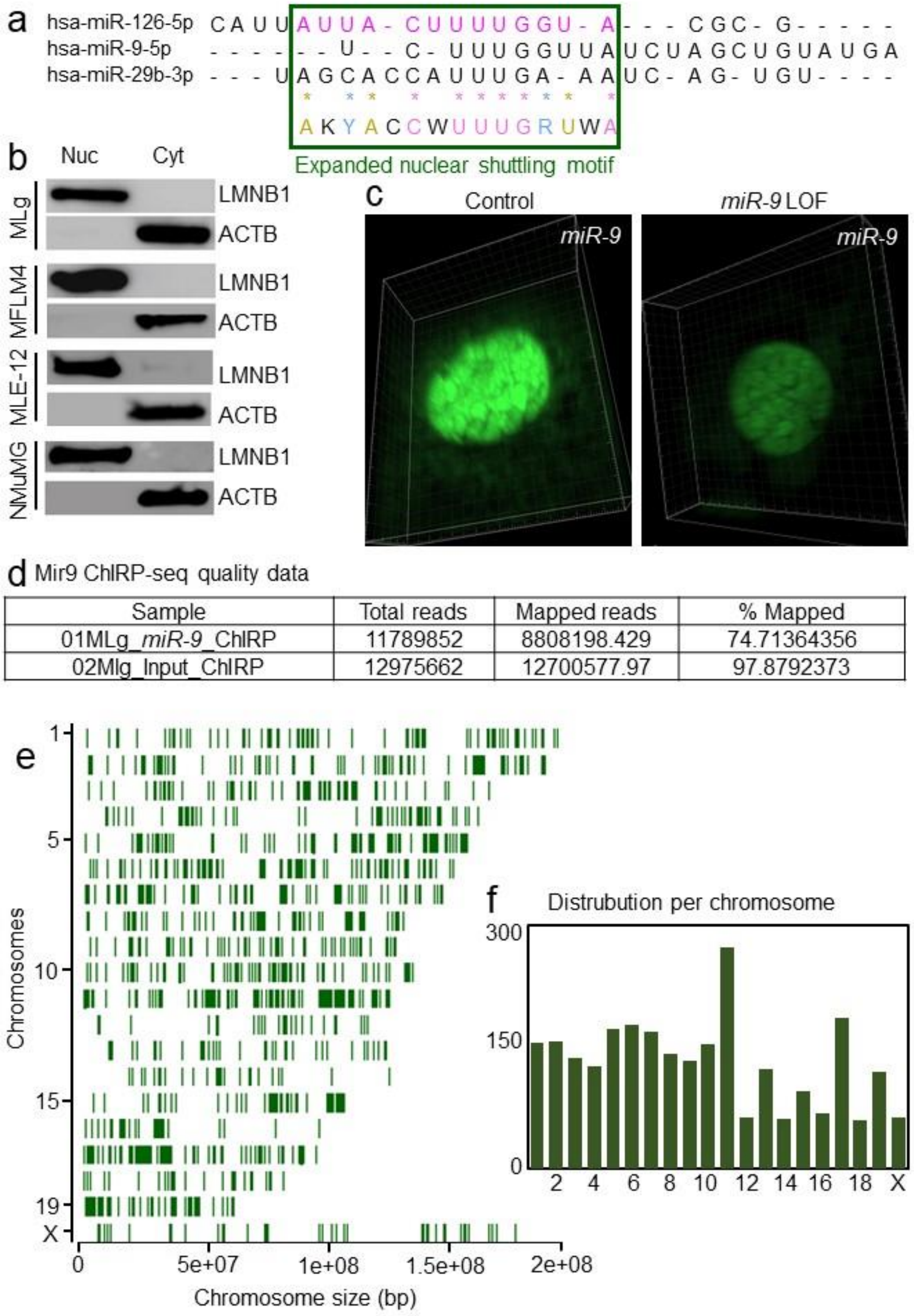


Supplementary Figure 1

**Supplementary Figure 1: Mature *miR-9* is detected in the cell nucleus at loci that are distributed on all chromosomes.** (**a**) Sequence alignment of the indicated mature human miRNAs highlighting the published *miR-126* nuclear shuttling motif (magenta) and the expanded miRNA nuclear shuttling motif (green square) using the IUPAC nucleotide code. The bottom line shows the partially conserved sequence. Pink letters are conserved among all sequences. Golden letters are conserved in at least 2 sequences. Blue letters indicate conserved type of base (either purine or pyrimidine). (**b**) Cell fractionation efficiency shown by Western Blot analysis of the nuclear (Nuc) and cytosolic (Cyt) fractions of the cell lines used for TaqMan assay-based expression analysis of mature *miR-9* in Figure 1b. LMNB1, lamin B1, nuclear marker; ACTB, beta actin, cytosolic marker. (**c**) Confocal microscopy of MLg cells after RNA FISH confirmed *miR-9* nuclear localization (left) and *miR-9* loss-of-function after *miR-9*-specific antagomiR probes transfection (right). (**d**) ChIRP-seq data sets supports the quality of the experiments. (**e**) Distribution of putative *miR-9* target loci on all chromosomes. (**f**) Number of putative *miR-9* target loci in each chromosome.

a H3K4me3 ChIP seq quality data

| Sample name | Total reads | Mapped reads | % Mapped |
|---|---|---|---|
| 1. Control | 44,049,013 | 42,309,773 | 96.05 |
| 2. *miR-9* LOF | 37,138,655 | 35,556,352 | 95.74 |

b

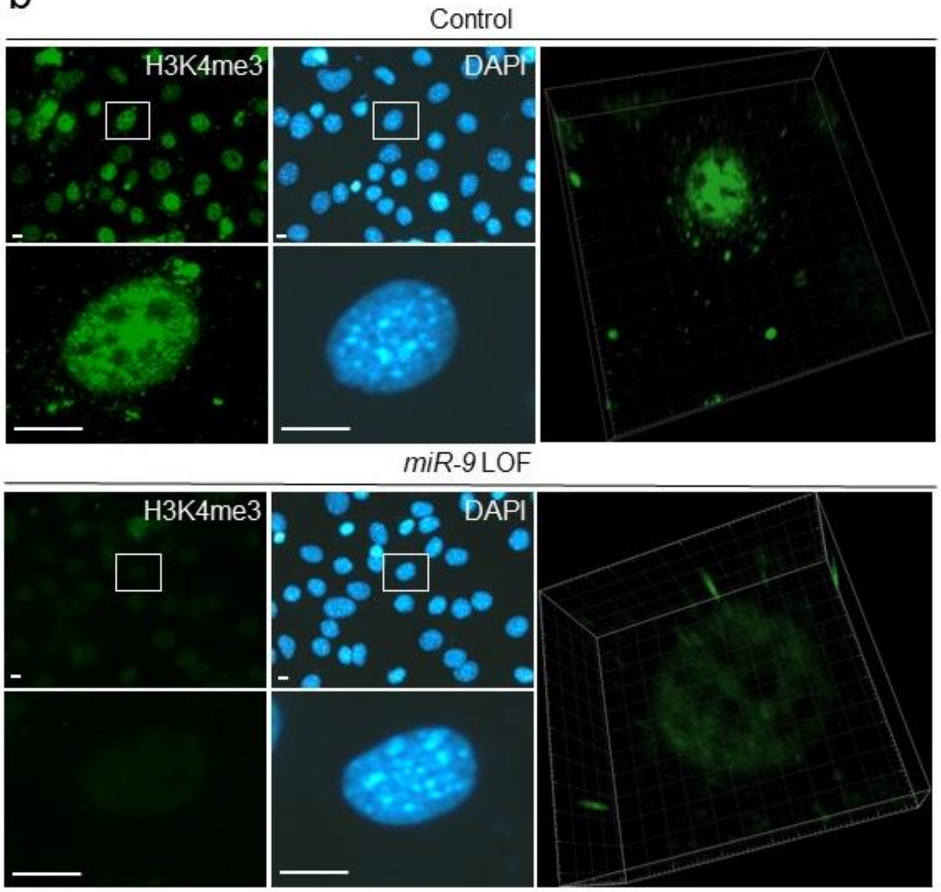


c RNA seq quality data

| Sample | Total reads | Mapped reads | % Mapped |
|---|---|---|---|
| 1. Control | 59,681,681 | 41,776,036 | 70 |
| 2. *miR-9* LOF | 50,055,390 | 36,778,286 | 73.48 |

Supplementary Figure 2

**Supplementary Figure 2: *MiR-9* is required for H3K4me3 broad domains** (**a**) Description of the ChIP-seq data set supports the quality of the experiment. (**b**) Fluorescence (left) and confocal (right) microscopy after H3K4me3-specific immunostaining of MLg cells transfected with control (top) or *miR-9*-specific (bottom) antagomiR probes to induce a *miR-9* loss-of-function (LOF) confirmed the reduction of H3K4me3 broad domains. Representative images from three independent experiments. Squares are shown at higher magnification at the bottom of the respective picture. Three-dimensional confocal images from single-cells (right). DAPI, nucleus. Scale bars, 10 μm. (**c**) Description of the RNA-seq data sets supports the quality of the experiment.

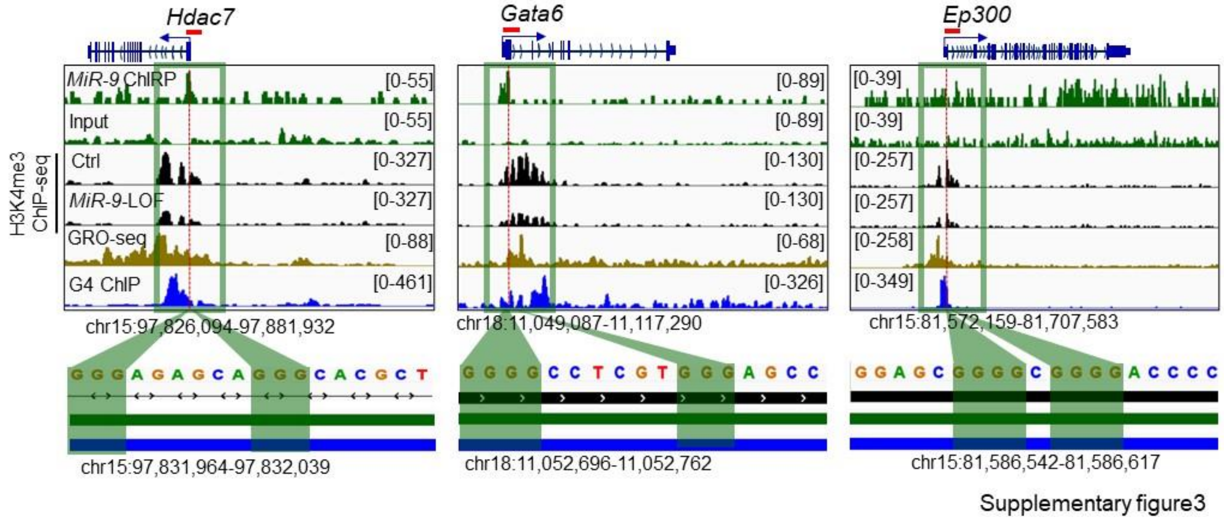


**Supplementary Figure 3: H3K4me3, nascent RNA and G4 are enriched at promoters of selected *miR-9* target genes.** Visualization of selected *miR-9* target genes (*Hdac7*, *Gata6* and *Ep300*) using IGV genome browser showing enrichment of *miR-9* by ChIRP-seq (green), H3K4me3 by ChIP-seq in Ctrl and *miR-9*-specifc antagomiR transfected MLg cells (black), nascent RNA by GRO-seq (brown) and G4 by G4P ChIP-seq (blue). Reads were normalized using reads per kilobase per million (RPKM) after bamCoverage. Images show the indicated gene loci with their genomic coordinates. Arrows, direction of the genes; blue boxes, exons; red lines, regions selected for single gene analysis; green squares, regions with enrichment of *miR*-9, H3K4me3, nascent RNA and G4; dotted lines, regions shown at the bottom with high G content.

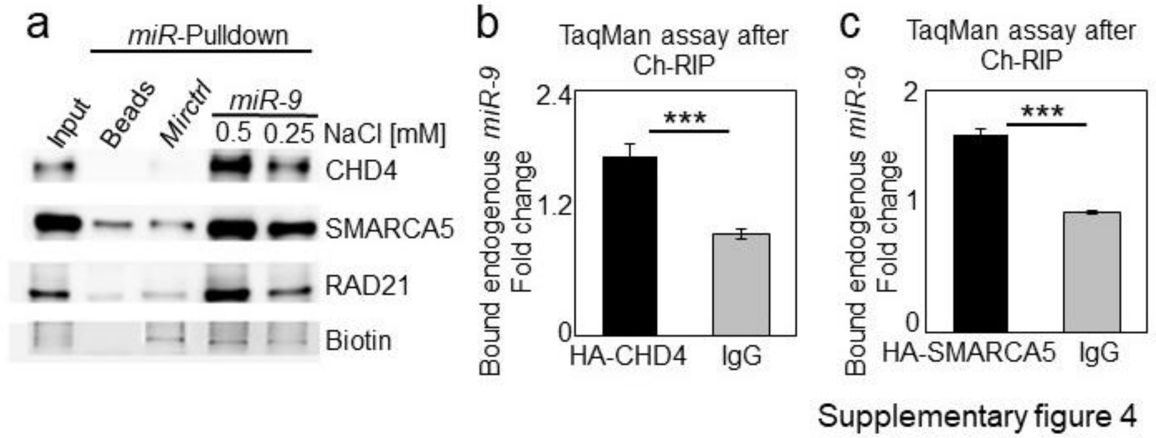


**Supplementary Figure 4: Nuclear *Mir9* interacts with CHD4, SMARCA5 and RAD21.** (**a**) Western blot of precipitated proteins after micro RNA pulldown (*miR*-Pulldown) using the nuclear fraction from MLE-12 cells transfected with biotinylated *Mirctrl* or *miR-9*. Input, 5% of starting material. Beads, negative control without biotinylated probe. *Mirctrl*, biotinylated scramble *miRNA*. *miR-9*, biotinylated *mmu-miR-9-5p*-RNA. NaCl in milimolar (mM). (**b-c**) Mature *miR-9*-specific TaqMan assays after chromatin RNA immunoprecipitation (Ch-RIP) using chromatin from MLE-12 cells transfected with expression constructs of HA-CHD4 (b) or HA-SMARCA5 (c) fusion proteins and HA-specific antibodies or IgG (negative control). Bar plots presenting fold change as means; error bars, s.e.m ($n$ = 3 biologically independent experiments); asterisks, $P$-values after two-tailed t-test, ***$P \leq 0.001$. Source data are provided as a Source Data file 01.

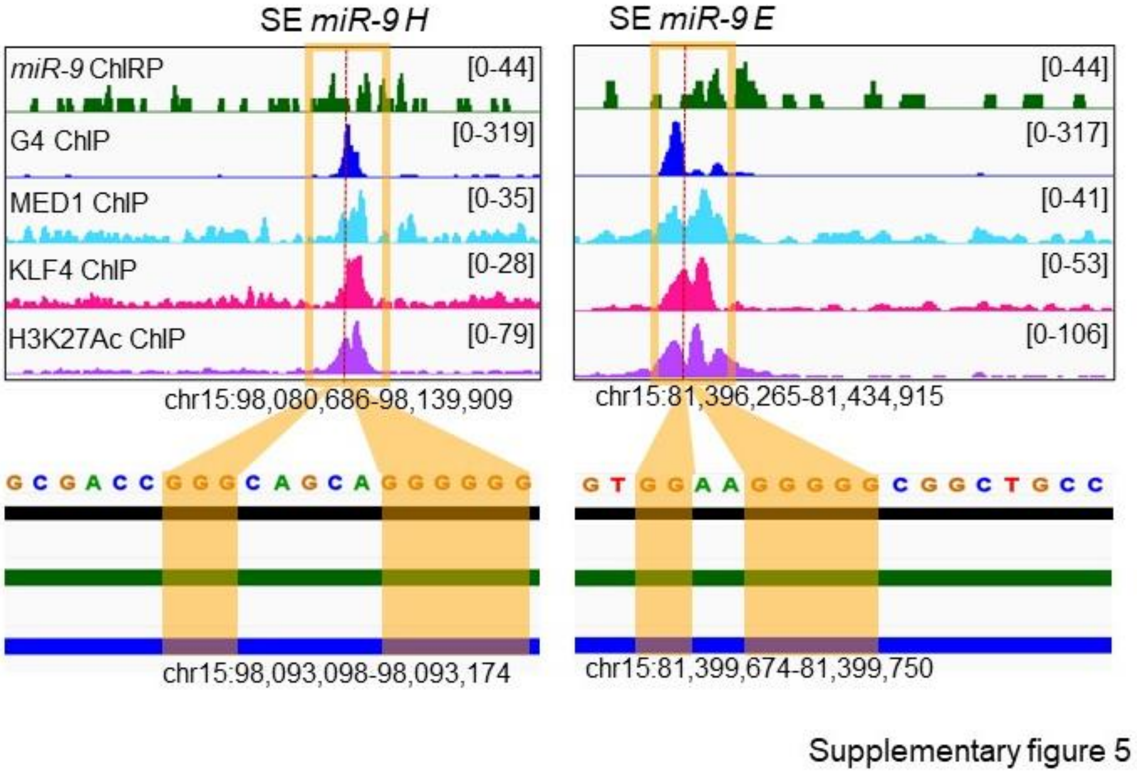


**Supplementary Figure 5: Nuclear *miR-9* is enriched at super-enhancers and is required for G-quadruplexes.** Visualization of selected SE (SE *miR-9 H* and SE *miR-9 E*) with *miR-9* enrichment using IGV genome browser showing enrichment *miR-9* by ChIRP-seq (green), G4 by G4P ChIP-seq (blue), MED1 (turquoise), KLF4 (magenta) and H3K27Ac (purple) by ChIP-seq. Reads were normalized using reads per kilobase per million (RPKM). Images show the indicated gene loci with their genomic coordinates. Orange squares, regions with enrichment of *miR-9*, G4 and SE markers; red lines, regions selected for single gene analysis; dotted lines, regions shown at the bottom with high G content.

a H3K4me3 ChIP seq quality data

| Sample name | Total reads | Mapped reads | % Mapped |
|---|---|---|---|
| 1. Control | 44,049,013 | 42,309,773 | 96.05 |
| 2.TGFB treated | 34,595,383 | 33,153,192 | 95.83 |
| 3. *miR-9* LOF + TGFB treated | 44,630,964 | 42,820,599 | 95.94 |

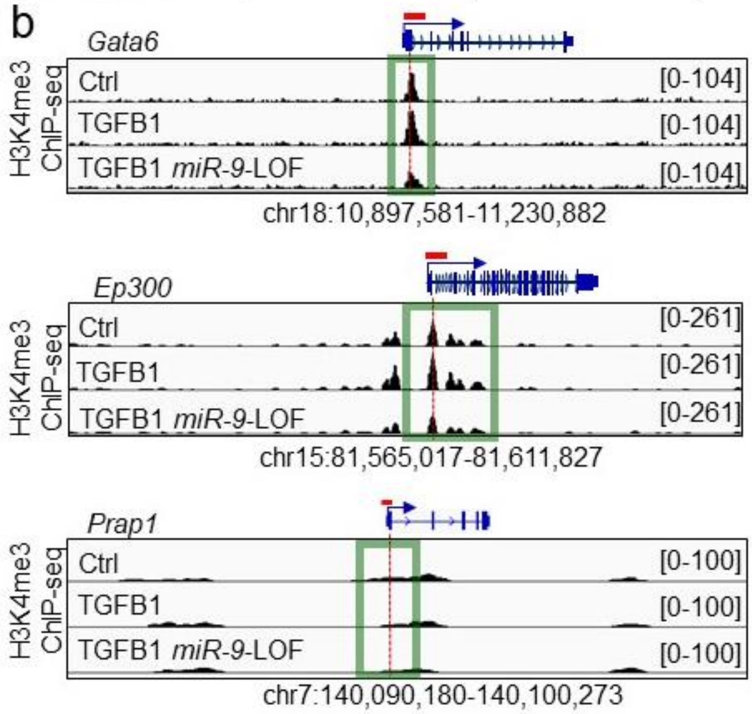


Supplementary figure 6

**Supplementary Figure 6: *miR-9* is required for H3K4me3 enrichment in the promoter region of TGFB1-responsive genes.** (**a**) H3K4me3 ChIP-seq data sets supports the quality of the experiments. (**b**) Visualization of *miR-9* target genes using IGV genome browser showing enrichment H3K4me3 by ChIP-seq in MLg cells that were transfected with control (Ctrl) or *miR-9*-specific antagomir to induce a loss-of-function (LOF), and non-treated or treated with TGFB1, as indicated. Images show the indicated loci with their genomic coordinates. Arrows, transcription direction; green squares, promoter regions; dotted lines, regions selected for single gene analysis.

a

| Sample name | Total reads | Full reads mapped | Trimmed reads mapped | Unmaped | % Mapped |
|---|---|---|---|---|---|
| 1. Control | 44049013 | 2840247 | 39469526 | 1739240 | 96.05 |
| 2.TGFB treated | 34595383 | 2010415 | 31142777 | 1442191 | 95.83 |
| 3.*MiR-9* LOF | 37138655 | 4144795 | 31411557 | 1582303 | 95.74 |
| 4.*MiR-9* LOF + TGFB treated | 44630964 | 2504104 | 40316495 | 1810365 | 95.94 |

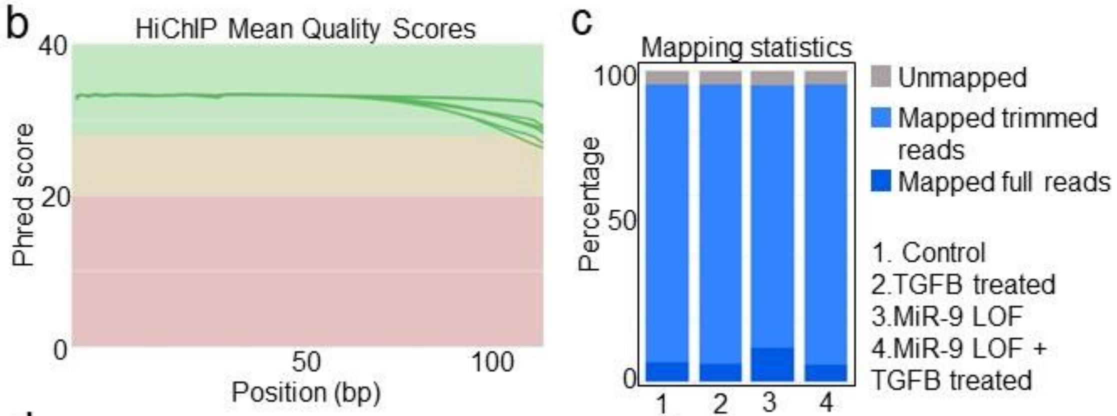


d

| Sample name | %Valid Pairs | %Invalid pairs) |
|---|---|---|
| 1. Control | 86.14 | 13.86 |
| 2.TGFB treated | 86.05 | 13.95 |
| 3.*MiR-9* LOF | 83.96 | 16.04 |
| 4.*MiR-9* LOF + TGFB treated | 87.56 | 12.44 |

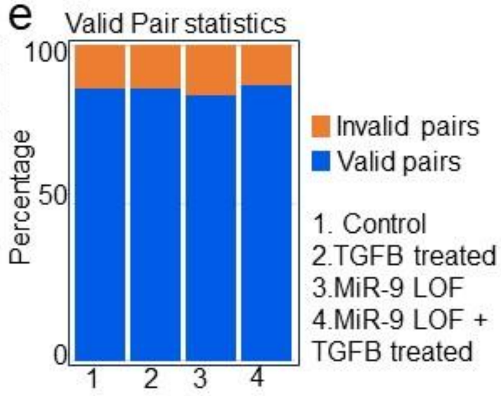


Supplementary figure 7

**Supplementary Figure 7: H3K4me3 HiChIP quality validation.** (**a**) Table showing the amount of reads, the trimmed, unmapped and the percentage of mapped reads. (b) Mean of Phred quality score. (**c**) Bar plot showing the distribution of the reads. (**d,e**) Pair distribution into Valid Pairs and Invalid Pairs, table (d) and bar plot (e).

**Data S1. (separate files)**

Source Data file 01 - This is an Excel file that contains the data for all the plots presented in the article, including the values for statistical significance and the implemented statistical tests.

Supplementary Table 1 – This is an Excel file that contains a list with the accession numbers of all published NGS data sets used in this manuscript.
